# DNA repair-based classification of melanoma cell lines reveals an effect of mutations in BRAF and NRAS driver genes on DNA repair capacity

**DOI:** 10.1101/2020.04.29.067900

**Authors:** Sylvie Sauvaigo, Manel Benkhiat, Florian Braisaz, Julien Girard, Sarah Libert, Stéphane Mouret, Florence de Fraipont, Caroline Aspord, Fanny Bouquet, Marie-Thérèse Leccia

## Abstract

Melanoma, the most serious form of skin cancer, frequently involves the dysregulation of key signaling pathways. Treatment strategies presently target the MAPK/ERK pathway, which is overactive in melanomas due in part to BRAF and NRAS mutations, and involve inhibitors against mutated BRAF (vemurafenib or dabrafenib) or MEK kinases (cobimetinib or trametinib), or a combination of the two. Using an established biochip technology, we assessed base excision repair (BER) and nucleotide excision repair (NER) activities in a collection of BRAF mutated (A-375, Colo 829, HT-144, Malme-3M, SK-mel5, SK-mel24 and SK-mel28) and NRAS mutated (M18, MZ2 and SK-mel2) melanoma cell lines, as well as wild-type controls (A7, CHL-1). We evaluated both basal activities (i.e., without treatment) and repair capacities after treatment with vemurafenib or cobimetinib alone, or in combination. Our results indicate that globally the DNA repair capacity of the cell lines was determined by the mutation status of the BRAF and NRAS genes, indicating that the MAPK pathway participates in the regulation of both BER and NER. Treatment of BRAF mutated melanoma cells with vemurafenib alone or the vemurafenib/cobimetinib combination, but not cobimetinib alone, led to reduced DNA repair capacity in about 60% of the BRAF mutated samples, indicating that signaling pathway inhibition can alter DNA repair activity. Upregulation of some DNA repair activities was also observed in several of the treated samples, suggesting activation of compensatory signaling pathways upon treatment. The data collectively indicate that mutations in the BRAF and NRAS genes exert distinct regulatory effects on the excision/synthesis steps of the BER and NER pathways and that targeted pharmacological inactivation of the signaling mechanism can translate into specific consequences in DNA repair capacity. The heterogeneity of the responses reported herein could help define subtypes of melanoma that are associated with resistance to targeted therapies.

## Introduction

Melanoma is the most dangerous form of skin cancer. It affects over 3 million individuals world-wide, killing roughly 60 thousand people annually, and exhibits dysregulation of several key signaling pathways. Mutations in the BRAF and NRAS proto-oncogenes play an early role in melanoma initiation and progression [1]. Around 50% of metastatic melanomas harbor *BRAF* mutations and 20% harbor an *NRAS* mutation, resulting in constitutive activation of the mitogen-activated protein kinase (MAPK) pathway [2].

The *BRAF* v600 point mutation is the most common oncogenic driver in melanoma and therefore constitutes a major molecular target for the development of therapeutic agents. The BRAF inhibitor vemurafenib (Vemu) was designed specifically to inhibit the MAPK (a.k.a., RAS-RAF-MEK-ERK) signaling pathway in tumors expressing the BRAF v600 mutant [3]. Mutated BRAF targeting agents produce an overall response rate above 50% in metastatic melanoma patients with a BRAF mutation, together with an improved progression-free and overall survival [3]. However, the success of BRAF inhibitors as a monotherapy has been compromised by the rapid onset of drug resistance, limiting the patients’ response to less than one year [3, 4].

Resistance to BRAF inhibitors is complex, with several mechanisms being involved, most of which result in reactivation of the MAPK pathway [5]. In addition, a high proportion of patients treated by anti-BRAF monotherapy require dose interruption or dose reduction due to toxicity. About 20% of patients experience proliferative and malignant cutaneous lesions attributed to paradoxical MAPK pathway activation or off-target kinase inhibition [6, 7].

The combined inhibition of BRAF and MEK (achieved with Vemu plus cobimetinib (Cobi)) in patients with BRAF V600 mutated metastatic melanoma is associated with a higher proportion of response and an increase in progression-free survival compared to anti-BRAF alone [7, 8]. However, as for BRAF inhibitor monotherapy, response to combined therapy is highly variable, the duration of the response differs greatly, and resistance emerges in most patients [5]. Reactivation of MAPK pathway is implicated, but activation of PI3K/AKT is also commonly observed through multiple mechanisms favoring survival and proliferation [9].

The molecular signaling in cancer is highly complex and offers redundancy, pathway cross-talk and feedback inhibition [10]. This intricacy makes it difficult to thoroughly understand the escape mechanisms that lead to resistance and the reactivation routes responsible for the paradoxical effects seen in melanoma patients. Consequently, intrinsic and acquired resistance to targeted therapies still represents a major challenge to treatment success, and the current molecular genetic diagnostics used to guide treatment strategy – i.e., *BRAF* and *NRAS* mutation detection - are not sufficient to predict tumor responsiveness and guide clinical approaches. For melanoma, several alternative genetic biomarkers (e.g., *NF-1, c-KIT, CDKN2A*) have been evaluated for their prognostic or predictive ability, but none have been validated so far or have proven clinical utility [5, 11]. Since treatment of patients with metastatic melanoma with checkpoint inhibitors only leads to a measurable response in a subset of patients, the identification of biomarkers that correlate well with positive outcome following treatment with targeted therapy remains critical.

DNA repair mechanisms and melanoma have long been associated. Ultraviolet (UV) irradiation is the most prominent recognized risk factor for melanoma [12, 13]. UVB (290-320 nm) and UVA (320-400 nm) contribute differently to UV-induced DNA lesions. Specifically, UVB radiation is directly absorbed by DNA and induces the formation of photoproducts, mainly cyclobutane pyrimidine dimers (CPDs) and pyrimidine-(6,4)-pyrimidone photoproducts (6-4PPs) [14]. UVA irradiation has a more complex form of action, as it has the capacity to generate not only photoproducts [15], but also reactive oxygen species that damage both DNA (e.g., 8-oxoguanine) and proteins that repair DNA damage [16]. DNA lesions generated by UVA and UVB may ultimately promote mutagenesis and skin carcinogenesis.

DNA excision repair systems [Nucleotide Excision Repair (NER) and Base Excision Repair (BER)] play an important role in removing UV-induced and oxidative lesions and preventing UV-related mutagenesis [17]. Moreover, DNA repair mechanisms have complex connections with the multiple regulatory signaling pathways involved in melanoma. For instance, the MAPK signaling pathway participates in the regulation of NER, and BER and the MAPK signaling pathway communicate through the RAS GTPase [18–20].

To gain further insights into the repair-signaling pathway relationship, we explored NER and BER mechanisms in melanoma cells mutated or not for the BRAF and NRAS genes. DNA repair capacities were evaluated at basal level (without treatment) or in response to BRAF and MEK inhibitor treatment alone or in combination. To that aim, we used a unique comprehensive strategy that investigated at a functional level, the repair of different oxidative and photoproduct DNA damage. Our goal was to identify biomarkers that could guide treatment decisions.

## Materials and Methods

### Cell Lines and Cell Culture

All melanoma cell lines were purchased from the American Type Culture Collection (ATCC), except two of the *NRAS* mutated cell lines (MZ2, M18), which came from other laboratories [21] and [22], respectively (Table 1). Cell culture products were from Gibco™. The cells were grown in RPMI-1640 medium supplemented with 10% Fetal Calf Serum (FCS), containing penicillin-streptomycin (50 U/mL and 0.05 mg/mL, respectively) and GlutaMAX 12 mM, at 37°C in a humidified 5% CO_2_ atmosphere. The cells were passaged when they reached 80% confluence. Cells were rinsed two times in PBS and detached using a Gibco™ trypsin-EDTA 0.05% solution. BRAF and NRAS genotyping was verified using a pyrosequencing method developed at CHU Grenoble Alpes and clinically employed.

**Table 1.** Cell lines characteristics and treatment applied (Cobi, Vemu or combination).

| Melanoma cell lines | Origin | Mutation | [Cobi] μM | [Vemu] μM | [Cobi+Vemu] μM |
| --- | --- | --- | --- | --- | --- |
| A7 | Skin | WT | 5 | 15 | 5 + 15 |
| CHL-1 | Skin | WT | 0.03 | 8 | 0.01 + 4 |
| A-375 | Skin | BRAFV600E | 0.005 | 0.3 | 0.003 + 0.1 |
| M18 | Skin | NRASQ61R | 0.03 | 8 | 0.01 + 4 |
| MZ2 | Skin | NRASQ61K | 0.03 | 8 | 0.01 + 4 |
| SK-mel2 | Skin | NRASQ61R | 0.03 | 8 | 0.01 + 4 |
| Colo 829 | Skin | BRAFV600E | 0.005 | 0.3 | 0.003 + 0.1 |
| HT-144 | Metastatic site: subcutaneous tissue | BRAFV600E | 0.005 | 0.3 | 0.003 + 0.1 |
| Malme-3M | Metastatic site: lung | BRAFV600E | 0.005 | 0.3 | 0.003 + 0.1 |
| SK-mel5 | Metastatic axillary node | BRAFV600E | 0.005 | 0.3 | 0.003 + 0.1 |
| SK-mel24 | Metastatic site: lymph node | BRAFV600E | 0.005 | 0.3 | 0.003 + 0.1 |
| SK-mel28 | Skin | BRAFV600E | 0.005 | 0.3 | 0.003 + 0.1 |

WT: wild-type for both NRA and BRAF S; NRAS and BRAF genes mutation are indicated. All were controlled by pyrosequencing.

### Cytotoxicity MTT Assay

Cells were seeded at a density of 3-6 × 10^4^ per well in 24-well plates (Falcon) in 450 µL of RPMI 1640 medium supplemented with 10% FCS and incubated at 37 °C for 24 h before treatments. Cobi (Genentech), Vemu (Genentech) and the combination were dissolved in dimethyl sulfoxide (DMSO, Sigma-Aldrich) as 10 mM stock solutions and used at different final concentrations (S1 Table and S1 Fig). Cells incubated with 0.1% DMSO served as controls for the cytotoxicity experiments. The MTT (3-(4,5-dimethylthiazol-2-y1)-2,5-diphenyl tetrazolium bromide) assay was used to assess cell viability as previously described in [23]. Cell viability was expressed as a percentage of the viability obtained for the controls (DMSO only). Data resulted from 3 independent experiments performed in triplicate.

### Cell Pellet Preparation

The concentrations of the inhibitors used in the DNA repair experiments were selected on the basis of cytotoxicity experiments. Specifically, the doses were selected because they induced about 20-30% toxicity (Table 1 and S1 Fig). Cells were treated for 24h and rinsed two times in PBS. Pellets containing 3-5 million cells were prepared and frozen at −80°C in albumin/DMSO (90/10, Sigma-Aldrich). Note that the wild-type for both BRAF and NRAS (WT) CHL-1 cell line appeared to be somewhat sensitive to Cobi. Three independent biological replicates were prepared for each of the selected experimental conditions.

### Nuclear Extract Preparation

Nuclear extracts were prepared as described earlier [24]. Frozen cell pellets were thawed and rinsed in PBS. They were then suspended in 1 mL Buffer A (10 mM KCl, 10 mM HEPES KOH, pH 7.8, 1.5 mM MgCl_2_, 0.02% Triton X-100) that contained 0.5 mM DTT (Sigma-Aldrich), 0.1 mM PMSF (phenylmethylsulfonyl fluoride, Roche) and incubated 10 min on ice. They were subsequently vortexed for 30s to complete the cytoplasmic membrane lysis. The nuclei were recovered by centrifugation for 5 min at 2300 g and suspended in 30 µL of Buffer B (400 mM KCl, 10 mM HEPES KOH, pH7.8, 1.5 mM MgCl_2_, 0.2 mM EDTA, 0.5 mM DTT, 25% glycerol, 0.1 mM PMSF, 0.7X antiprotease (Complete-mini, Roche)). After a 20 min incubation on ice, the nuclear membrane lysis was completed by 2 cycles of freezing-thawing in liquid nitrogen – 4°C respectively. Membrane debris were cleared by centrifugation for 10 min at 16 000 g. The supernatant was stored as aliquots at −80°C until used. The protein content of the cell free extracts was assessed using the Uptima MicroBCA Protein Quantification kit (Interchim). Typical protein concentration was 2-4 mg/mL.

### DNA Repair Assays

The excision/synthesis DNA repair capacities of the samples were determined using the ExSy-SPOT assay (LXRepair, Grenoble, France). The assay has been extensively described [25, 26]. Briefly nuclear extracts were diluted to a final protein concentration of 0.3 mg/mL in Repair Buffer (1X composition: 6 mM Hepes-KOH pH 7.8, 80 mM KCl, 7.3 mM MgCl_2_, 2 mM EDTA-NaOH pH 8.0, 0.6 mM DTT, 8.4% glycerol) containing 0.25 μM of each of the unlabeled-dNTPs (dA, dT, dG), 10 mM phosphocreatine (Sigma-Aldrich), 50 μg/mL creatine phosphokinase (Sigma-Aldrich), 0.1 mg/mL BSA, 1 mM ATP and 0.25 µM dCTP-Cy3 (GE healthcare). The mixtures were then applied to biochips functionalized with a series of plasmids containing specific lesions. Five substrates allowing the characterization of five different repair activities were available on the biochip: 8oxoG (8-oxo-dGuo, repaired by BER and involving OGG1), AbaS (abasic sites, repaired by APE1), CPD-64 (photoproducts: CPDs and 6-4PPs, repaired by NER), Etheno (Etheno adducts, repaired by BER and involving MPG [27]), and Glycols (pyrimidine Glycols, repaired by NTH1 that belongs to BER [28]). In addition all investigated repair pathways require polymerase (pol) β or polδ/ε for DNA synthesis. A lesion-free plasmid was immobilized in parallel and served as a control. Repair of the substrates on the biochip by the DNA repair enzymes contained in the extracts led to the incorporation of the fluorescent dCTP. The repair reactions were run on 2 separate biochips for 3h at 37°C. The biochips were then washed 2×5 min in PBS, rinsed in H_2_O, and dried 10 min at 37°C. Images were then acquired at 532 nm using a Innoscan 710 AL scanner (Innopsys, Toulouse). Total spot fluorescence intensity (FI) was quantified using the Mapix 6.5 software (Innopsys). Data collected from the replicates were normalized using NormalizeIt software (LXRepair). The level of fluorescence incorporated into the lesion-free plasmid was subtracted from the fluorescence incorporated into each lesion-plasmid. A mean FI (+/− Standard Deviation) was calculated from the 3 replicates for each repair pathway.

Each sample, in each treatment condition, was thus characterized by a set of 5 FI values corresponding to the repair efficacy of the 5 repair pathways investigated. This set constituted a highly specific DNA Repair profile with simultaneous quantitative information on the repair efficacy of each of the repair mechanisms analyzed.

### Statistical Analysis

For the box-plot representation and the classifications, values of the whole dataset were standardized (mean = 0 and standard deviation = 1). To analyze the influence of the mutation on the DNA Repair signature, we used the non-parametric Kruskal-Wallis test. We considered that this influence was significant between the mutation groups when p<0.05. In that latter case, we further applied the post hoc non parametric Nemenyi test, to more precisely characterize the differences for each repair pathway, by testing all the possible pairwise combinations. The effect of the compounds alone and in combination on the different DNA repair activities was examined using the value of ratio of the FI obtained for the repair of each lesion in each treatment condition (T) versus the non-treated condition (NT) (R= FI T/ FI NT), independently for each of the triplicates. A ratio of 1 means that the compounds have no activity, if R<1 that means that the treatment inhibits DNA repair activities and R> 1 indicates an increase in the repair activities. We then determined if this ratio was statistically different from 1 using the Student t test.

### Data Presentation

#### Box-Plot

The data were displayed as box-plots to visualize 1. the qualitative and quantitative consequences of the BRAF and NRAS mutations on the DNA repair profiles, and 2. the impact of the treatments according to the mutation status. The red line inside the box represented the median. The first and third quartiles were identified by the box limits, and minimum and maximum by the external bars. The outliers were plotted as individual + symbol.

#### Classification

To better visualize possible DNA repair capacity dissimilarities between mutation groups, we clustered the cells lines according to their specific DNA repair profiles, using unsupervised hierarchical clustering (unweighted pair group method with arithmetic mean (UPGMA)), with the software R (http://r-project.org/). The hierarchical average linkage clustering algorithm was run using the Euclidean distance, which aggregates profiles with similar FI levels and covariation. The optimal number of clusters was determined by cutting the cluster dendrograms at the agglomeration criteria inflexion point. Clusters with an AU p value > 95% were considered significant.

Within each cluster, the v-test was used for ranking the variables according to their influence (S2 Table).

## Results

### DNA repair capacities without or with Cobi and/or Vemu treatment

To explore the consequences of mutations in NRAS and BRAF genes on DNA repair and to determine the impact of treatment on NER and BER capacity according to mutation groups (WT, NRAS mutated and BRAF mutated) in a collection of melanoma cell lines (Table 1), we employed a functional multiplexed assay on a biochip that allows a comprehensive and quantitative evaluation of repair mechanisms for a set of different DNA substrate lesions (BER: 8oxoG, AP sites, ethenobases, glycols; NER: CPDs and 6-4PPs). In brief, the assay measures the excision/synthesis capacities of DNA repair enzymes contained in lysates prepared from the samples of interest. The level of incorporation of fluorescent dNTPs during the synthesis step that occurs after the excision of the lesions reflects the global DNA repair capacity of the lesion-specific pathways.

Cells (Table 1) were either treated or not with Cobi, Vemu, or its combination (Cobi+Vemu), followed by nuclear extract preparation and DNA repair capacity measurement. The data (FI for repair of each of the 5 lesions) obtained from 3 independent experimental runs were pooled for the analysis, and significance was determined using the non-parametric Nemenyi test. The collective results are shown as Box-Plots for each repair pathway separately by mutation group and by treatment (Fig 1), representing the capacity of the indicated cells to repair each lesion.

**Fig 1.**
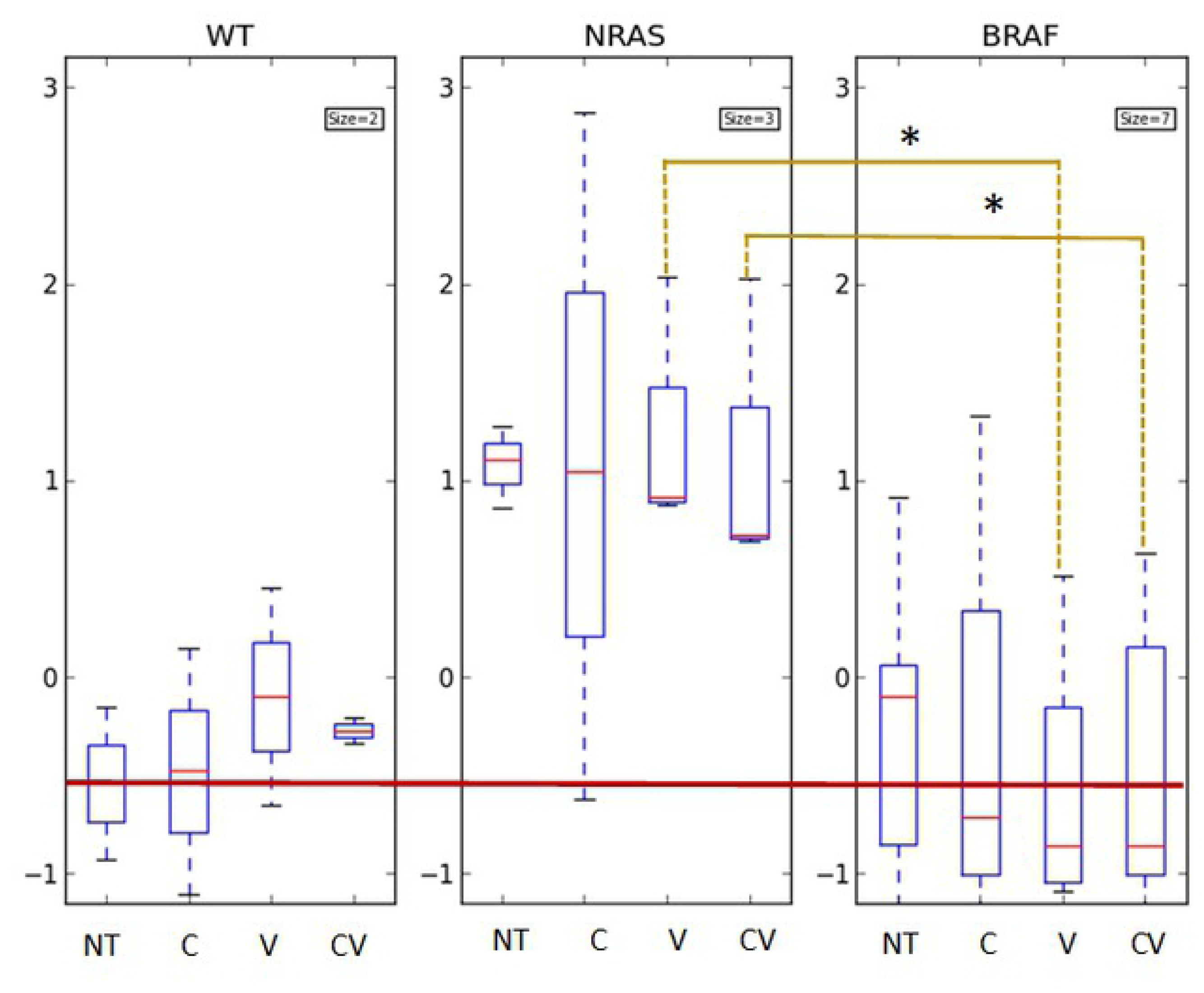
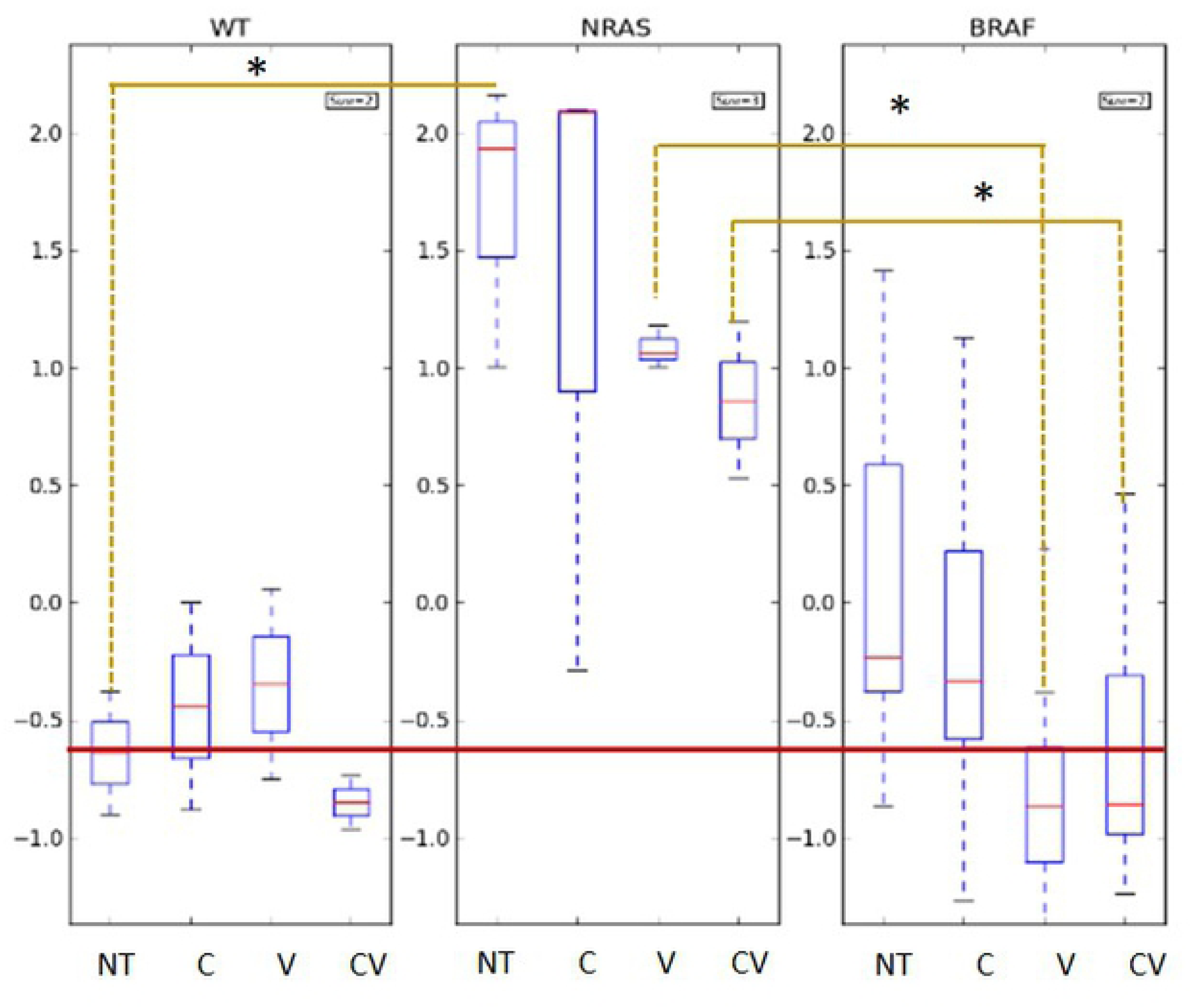
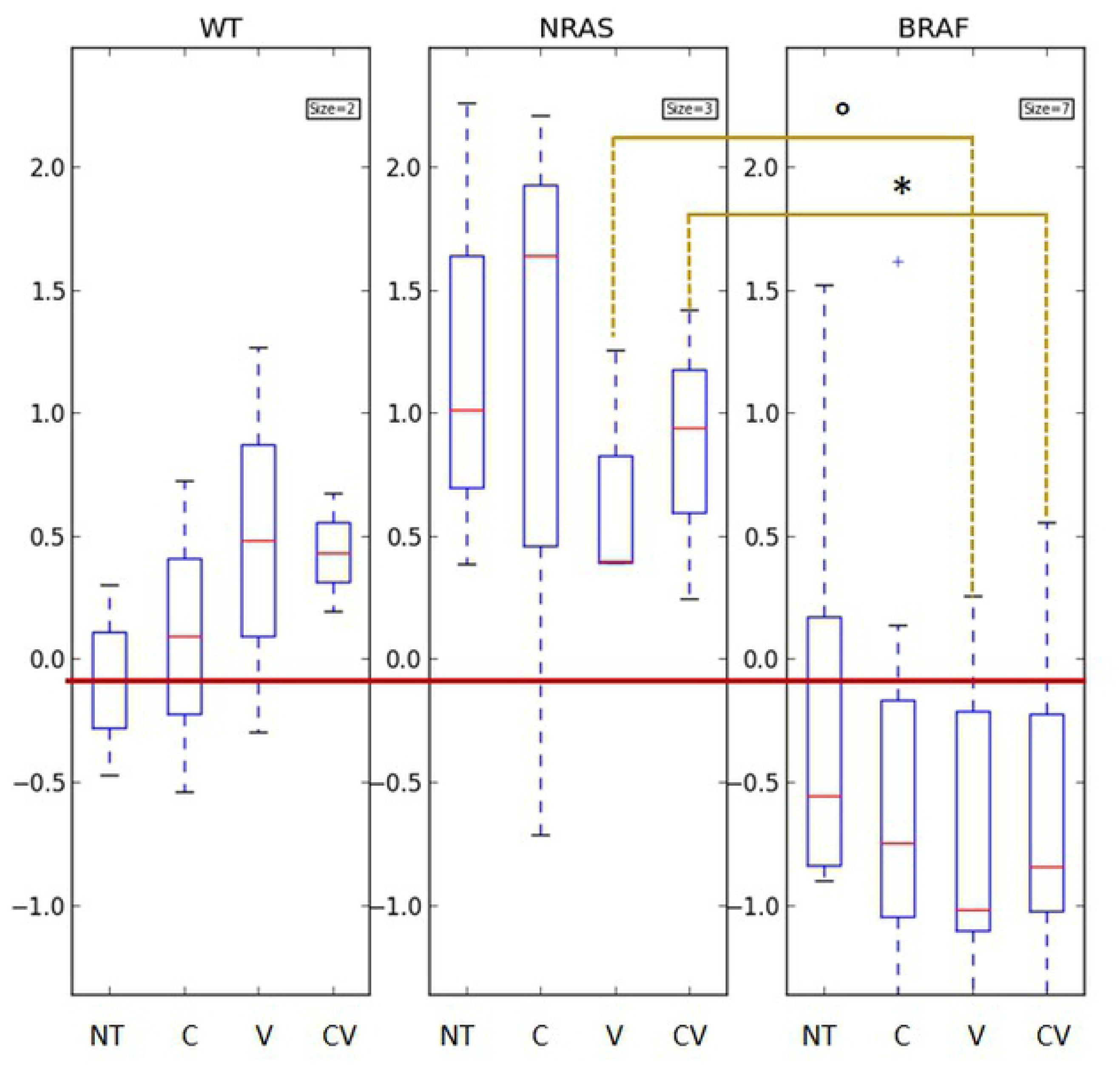
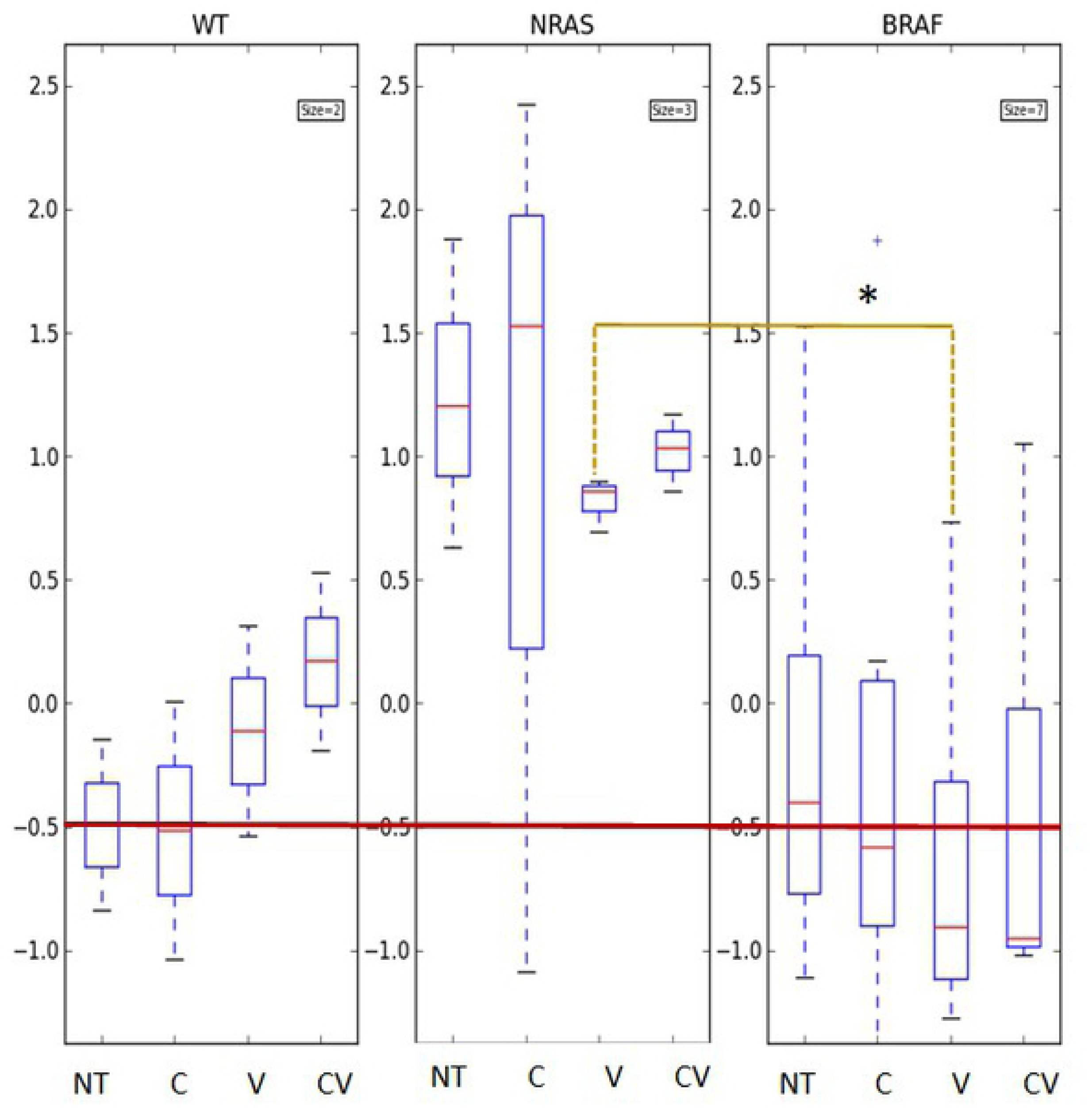
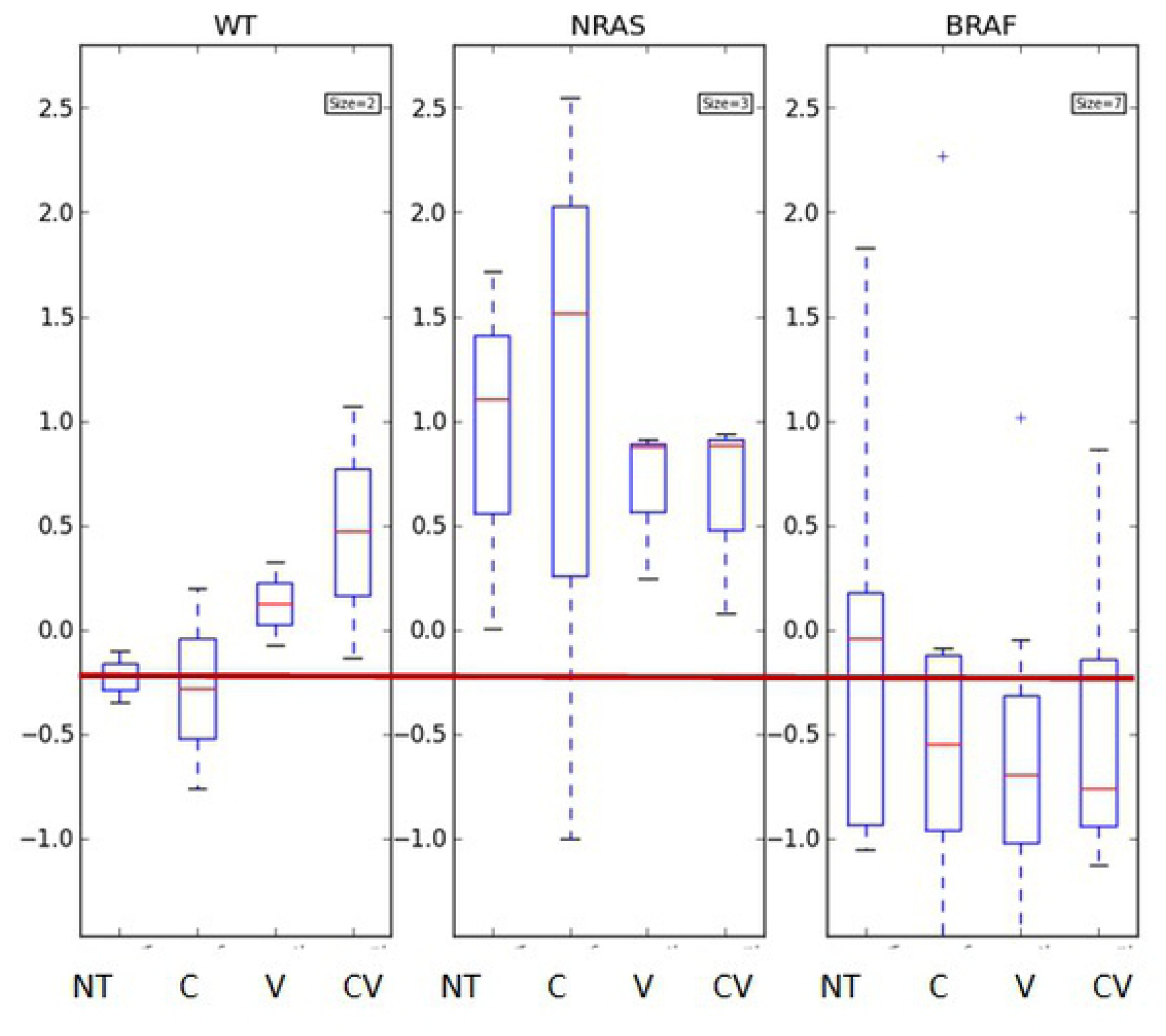
DNA repair capacities of the cell lines according to mutations in BRAF and NRAS genes, and consequences for the impact of therapeutic agent treatment on DNA repair capacity. Box-Plots displaying the DNA repair capacities of each repair pathway (for the following lesions: **A.** 8oxoG, **B.** AP sites, **C.** Photoproducts (CPDs and 6-4PPs), **D.** Ethenobases, **E.** Glycols) for each group of cells (WT, mutated for NRAS, mutated for BRAF). Data obtained in non-treated conditions (NT) or following treatment by Cobi (C), Vemu (V) and Cobi+Vemu (CV) are shown for the 3 groups of cells. Data expressed as FI were standardized. The red line within each box indicates the FI median value of each group.

The red line across the 3 diagrams, positioned at the level of the median value of the first displayed result (NT for WT cell lines), is a visual reference to help compare the results between groups and the impact of the treatment.

A. Repair of 8oxoG
B. Repair of AP sites (AbaS)
C. Repair of photoproducts (CPD-64)
D. Repair of ethenobases
E. Repair of glycols

Differences between groups: * p< 0.05; ° p = 0.056

In the absence of treatment, there was a trend towards lower DNA repair capacity in the WT and BRAF mutated groups compared to the NRAS mutated group. However, it was significant (*p*-value = 0.035) only when comparing the NRAS and WT groups for the repair of AP sites (AbaS, Fig 1).

Notably, differences in DNA repair capacity between the NRAS and BRAF groups reached significance when the cells were treated with Vemu or Cobi+Vemu, but not with Cobi alone (Fig 1), meaning that only treatment regimens involving Vemu modified the DNA repair capacities among the mutation groups. In all situations involving a significant change in DNA repair capacity, repair was higher in the NRAS group compared to the BRAF group. These treatment-specific differences were observed for the 8-oxoG and AbaS substrates, CPD-64 (Cobi+Vemu only), and Etheno (Vemu only) (Fig 1). It is worth mentioning, although the results did not reach significance, that repair of AP sites in the NRAS group appeared affected by Vemu and Cobi+Vemu.

### Comparison of DNA repair signatures across the different cell lines and treatments using hierarchical clustering

Hierarchical clustering is a convenient analysis tool to identify similarities between the elements of a group and to identify specific subgroups with distinct features. In order to determine whether cells with identical mutations clustered together, when untreated on the one hand, and after treatment on the other hand, we analyzed the DNA repair profiles obtained in the different conditions using hierarchical classification. At basal level (i.e., untreated, Fig 2A), four classes of DNA repair signatures were identified. Three cell lines, all belonging to BRAF group, displayed very low DNA repair capacities (Class 1_NT): Colo-829, A375 and SK-mel24 (defined as the “low repair” class; significant for all repair activities (S2 Table). Notably, they were clustered with one of the WT cell lines (CHL-1). In contrast, SK-mel28, SK-mel2 and M18 exhibited the most elevated DNA repair capacities (Class 4_NT; significant for all repair activities), with one cell line belonging to the BRAF group and the other two belonging to NRAS group. In between these 2 Classes were Class 2_NT, which includes 3 BRAF (SK-mel5, HT-144, Malme-3M) and 1 WT (A7) cell lines, and the NRAS cell line (MZ-2) alone (Class 3_NT). Notably, most BRAF cell lines had comparatively low DNA repair capacities (low FI signals), whereas most NRAS cell lines displayed high DNA repair capacities (high FI signals).

**Fig 2.**
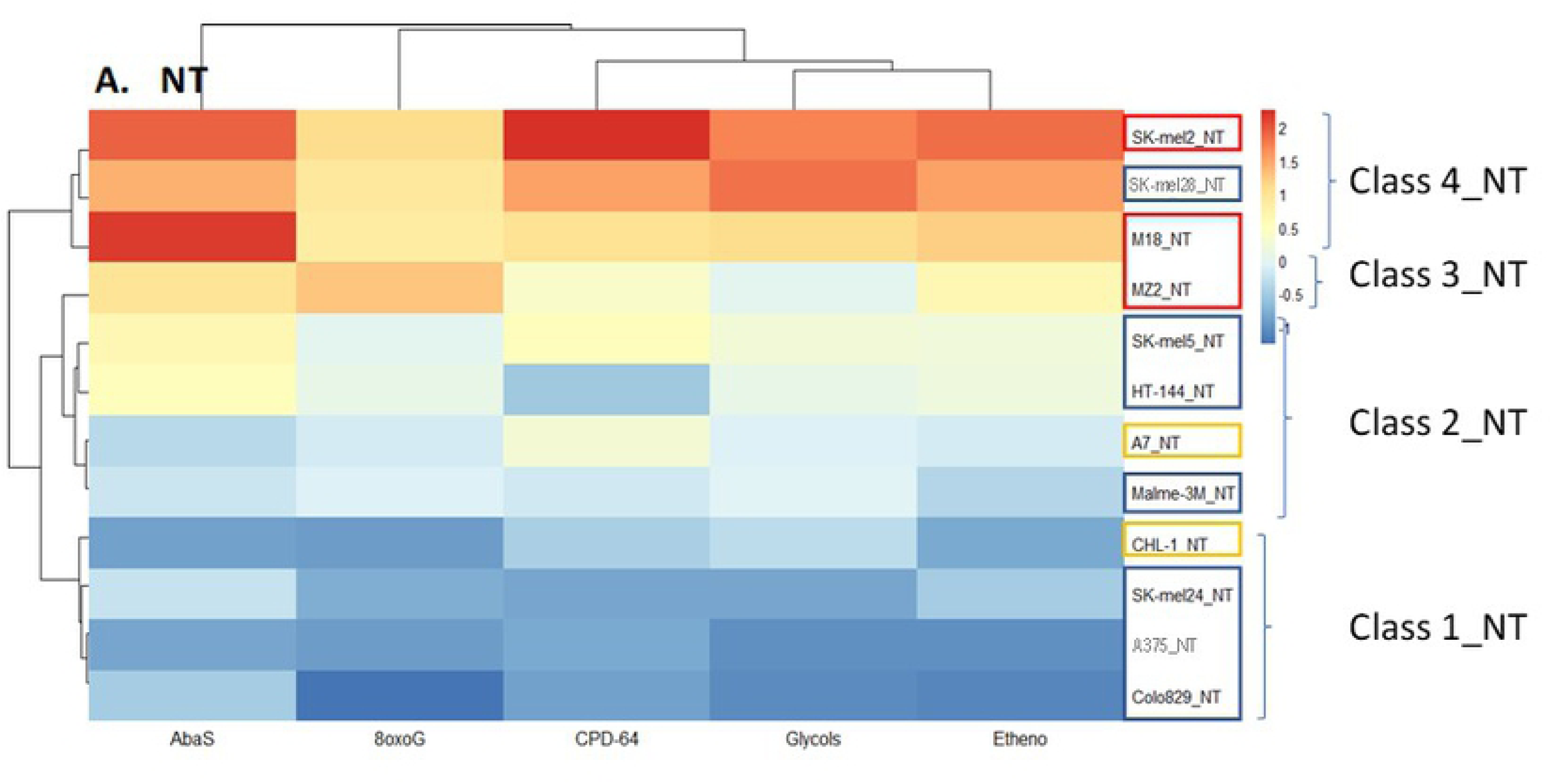
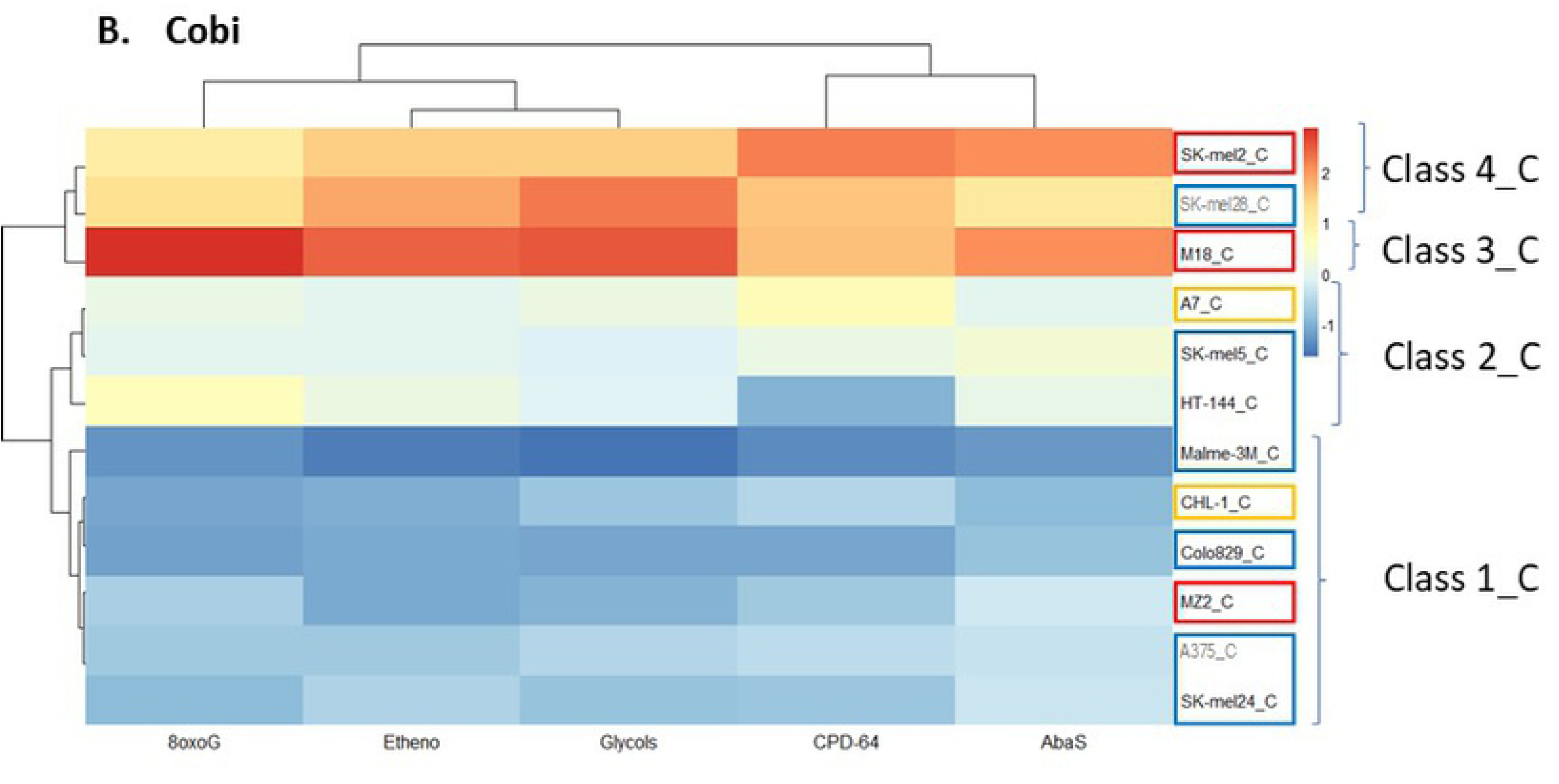
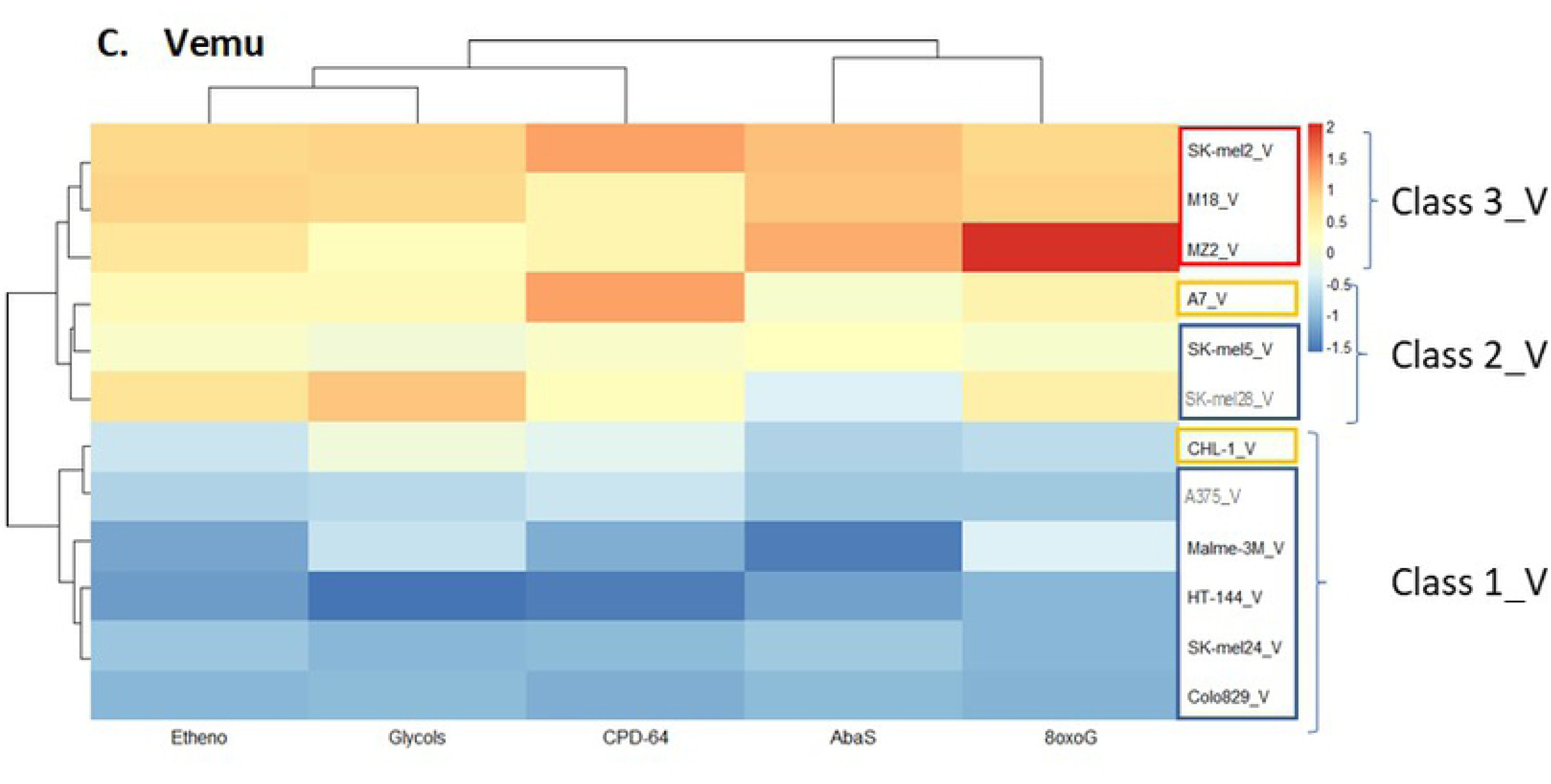
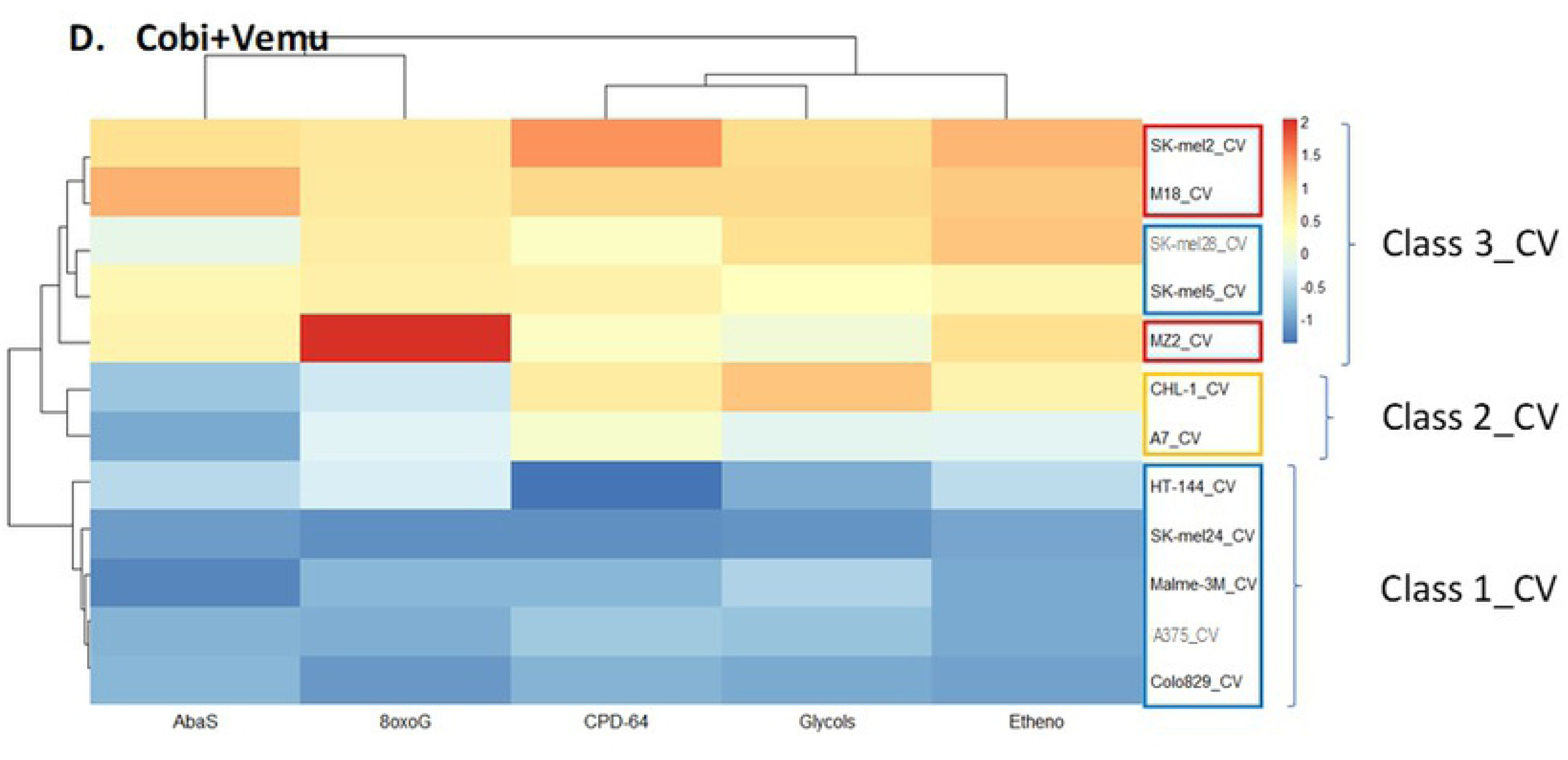
Comparison of the DNA repair capacity of the different cell lines according to their DNA repair signatures (hierarchical clustering). The heat maps represent the repair capacity of each pathway, represented by the lesion (substrate) it handles (8oxoG, AbaS, CPD-64, Etheno, Glycols) across cell lines. Data were normalized across the dataset: white represents the mean, red represents activity above mean, and blue represents activity below the mean **A.** Results obtained in non-treated (NT) conditions. **B**. Results obtained when cells were treated by Cobi. **C.** Results obtained when the cells were treated by Vemu (V). **D.** Results obtained when cells were treated by Cobi+Vemu (CV). Clustering revealed several distinct classes indicated by the numbers (See S2 Fig for details). DNA repair substrates are indicated below the figures, cell lines are indicated on the right of the figure. WT cell lines are framed by yellow rectangles, NRAS cell lines are framed by red rectangles and BRAF cell lines are framed by blue rectangles.

To next investigate the impact of the therapeutic compounds on DNA repair activities and to compare the results between the mutation groups, we clustered the DNA profiles obtained after treatment, independently for each condition (Cobi, Vemu, Cobi+Vemu; Figs 2B, 2C, 2D respectively). In experiments involving Cobi treatment alone (Fig 2B), the main change compared to the untreated condition was a shift of the NRAS mutated MZ2 cell line towards the low repair group (Class 1_C of Fig 2B). This observation shows that treatment with Cobi had a negative impact on the DNA repair capacities of the MZ2 cell line specifically. The other clusters remained almost unchanged, indicating that the DNA repair capacities of the other cell lines were globally unaltered by this therapeutic agent.

Treatment with Vemu had a greater impact on the hierarchical classifications, with three classes being distinguishable (Fig 2C). Cell lines with the lowest DNA repair activities (Class 1_V) consisted of 5 BRAF cell lines (the “low repair” class (Class 1_NT: Colo-829, A375, SK-mel24), joined now by Malme-3M and HT-144) and WT CHL-1. All DNA repair activities were significantly down regulated within Class 1_V (S2 Table). The 3 NRAS cell lines (SK-mel2, M18 and MZ2) that exhibited the highest repair activities when untreated retained high repair capacities when treated by Vemu and clustered together (Class 3_V; significant for 8oxoG, AbaS and Etheno (S2 Table)). Notably, treatment by Vemu completely separated NRAS cell lines from the other cell lines in classification, showing, as expected, that Vemu had a more specific impact on BRAF mutated cells, while NRAS mutated cells, which were not impacted, remained apart. Class 2_V, with intermediate DNA repair activities, included the remaining BRAF cell lines (SK-mel28, SK-mel5) and one of the WT cell lines (A7).

Similar to the classification obtained for the Vemu treated cells, the high repair capacity class following Cobi+Vemu treatment (Fig 2D) consisted of the 3 NRAS cell lines (Class 3_CV; SK-mel2, M18, MZ2; significant for all repair activities). However, with the combined treatment, the BRAF cell lines, SK-mel28 and SK-mel5, also clustered within the high repair capacity Class 3_CV. The “low repair” BRAF mutated group identified with Vemu treatment remained unchanged (Class 1_CV, encompassing Colo-829, Malme-3M, SK-mel24, A375 and HT-144), and the 2 WT cell lines made up an independent, intermediate group within the Cobi+Vemu collection (Class 2_CV) (S2 Table).

### Specific effect of the treatments on each cell line analyzed through FI treated/untreated ratios

As shown in Fig 2, cell lines from a same mutation group could behave differently with respect to their response to the different treatments. Indeed, several patterns of responses were observed. To better describe this variability, we aimed to gain insight into each cell line response to the treatments individually. Towards that goal, we calculated the ratio of FI treated to FI untreated (FI T/FI NT) for each repair activity, each cell line, under each treatment condition. The resulting histograms are displayed in Fig 3. We then used the Student-t test to determine which ratios (calculated from 3 independent biological replicates) were significantly different from 1 (based on p-values <0.05), considering each repair activity separately.

**Fig 3.**
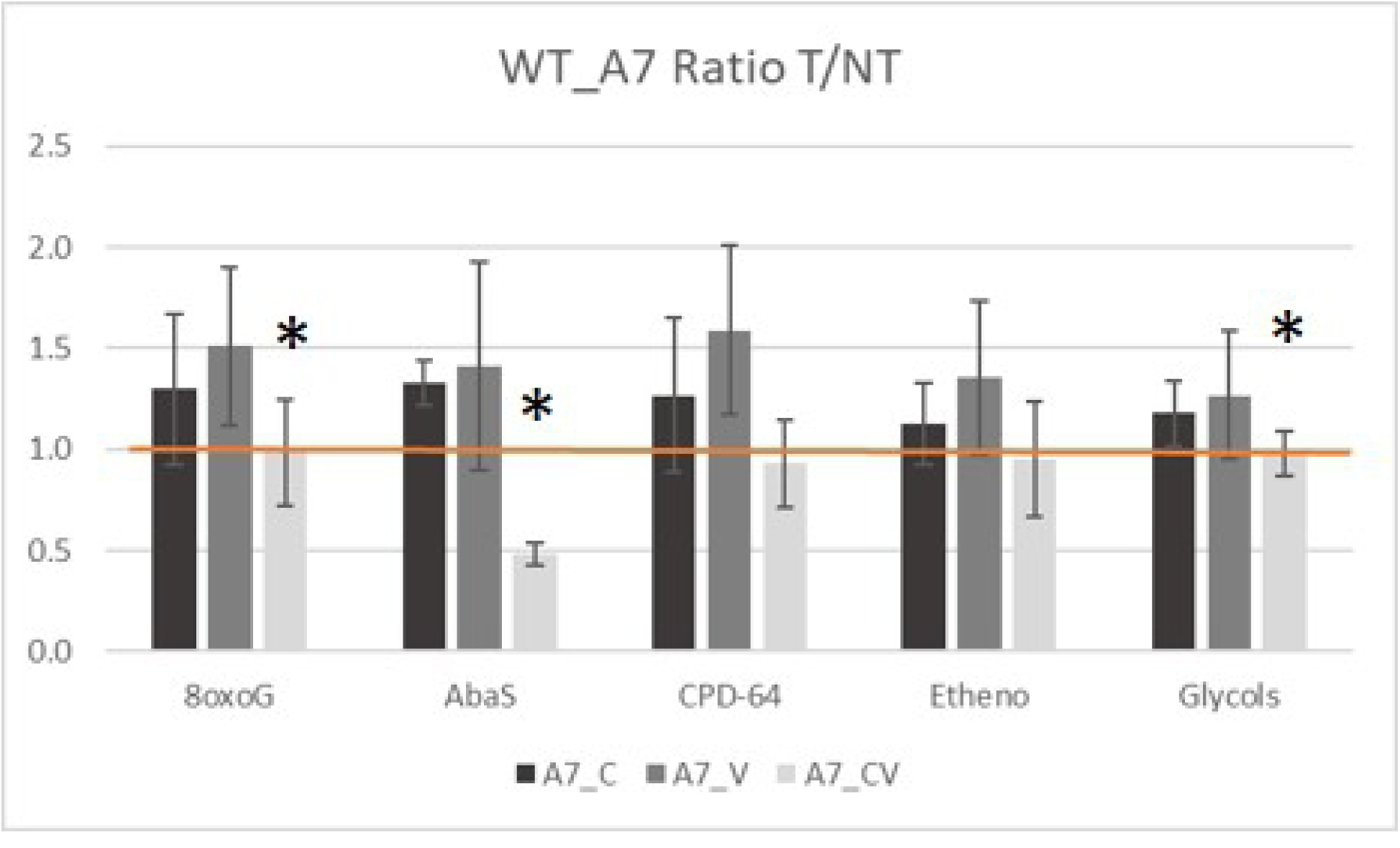
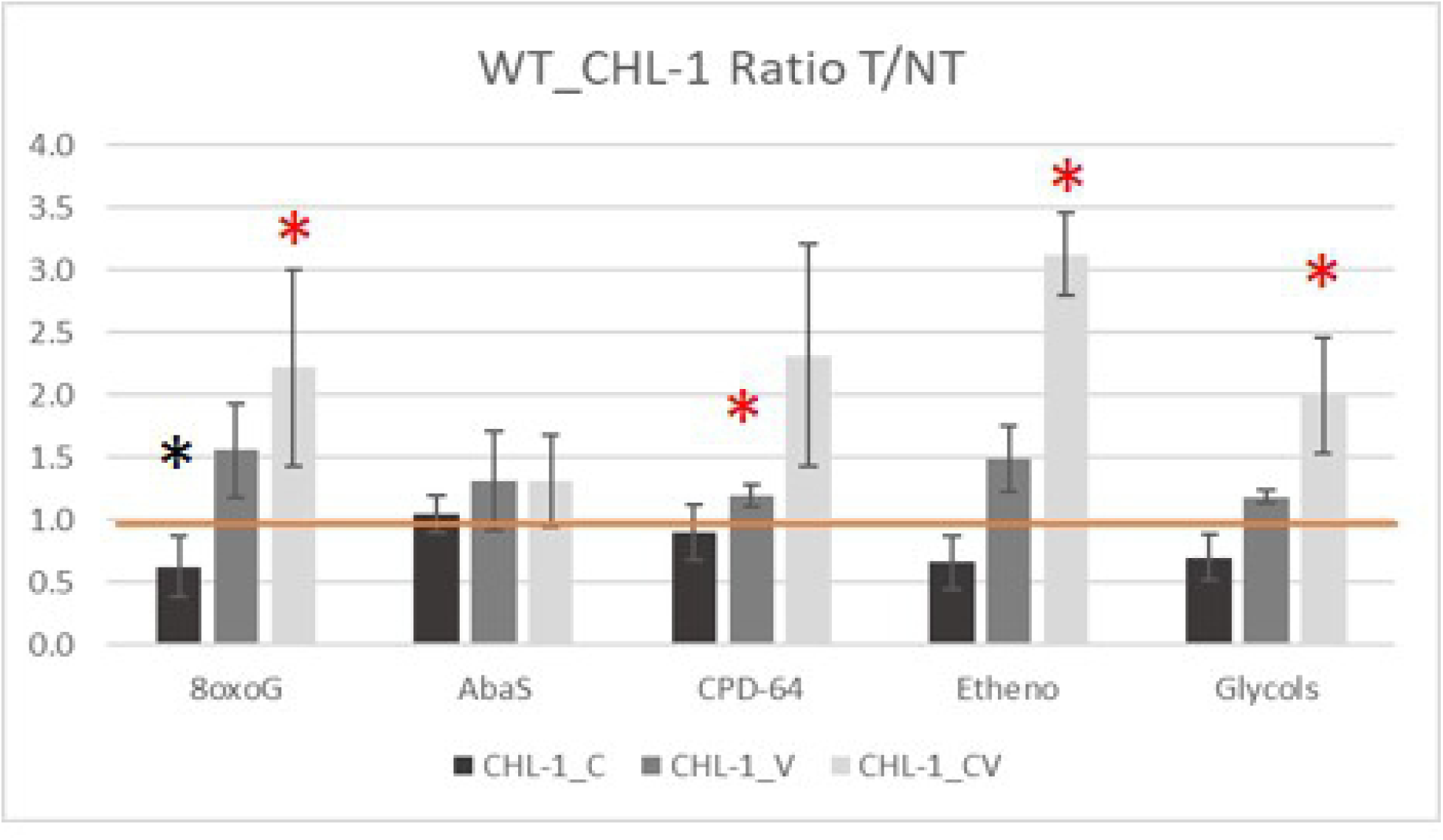
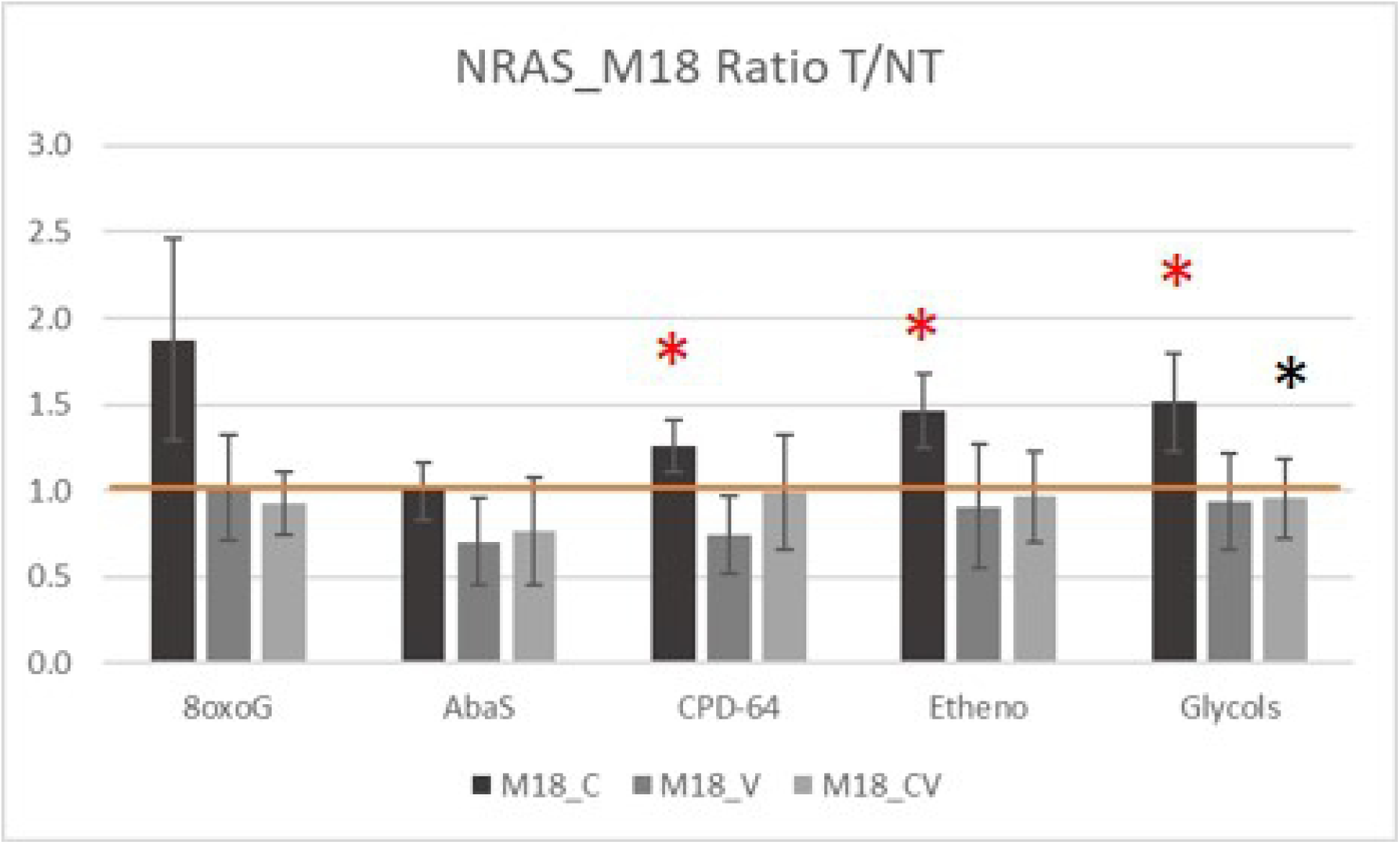
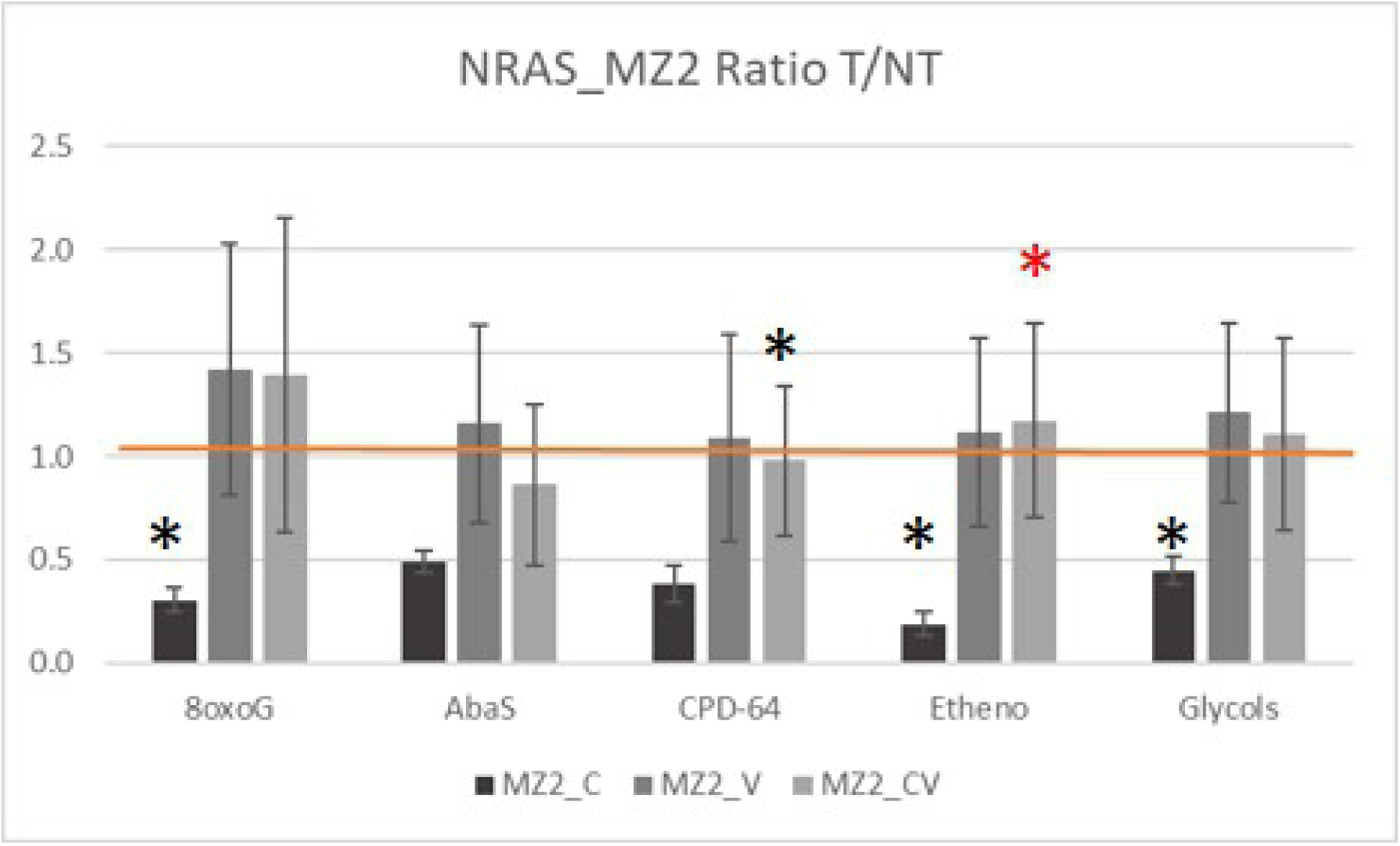
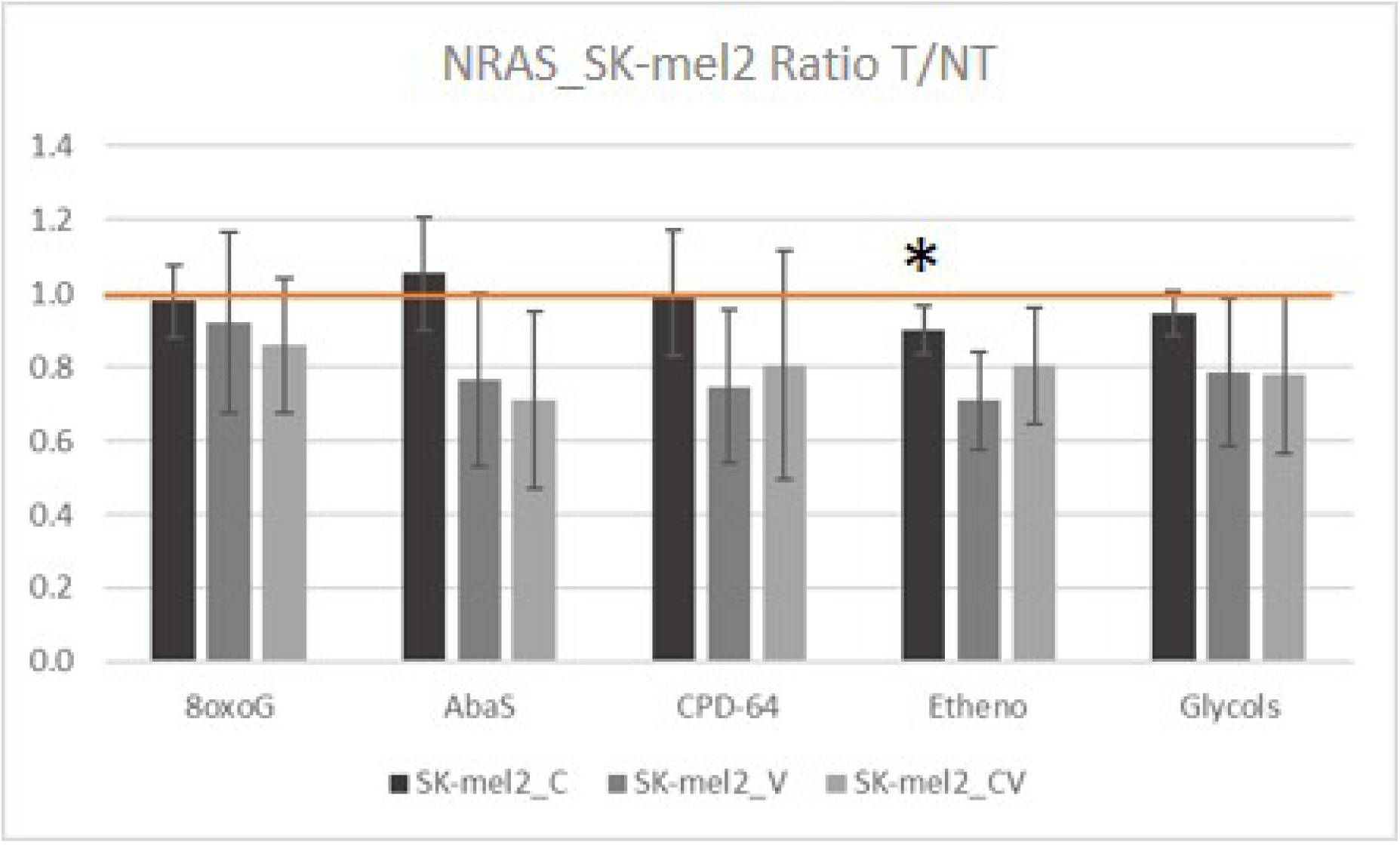
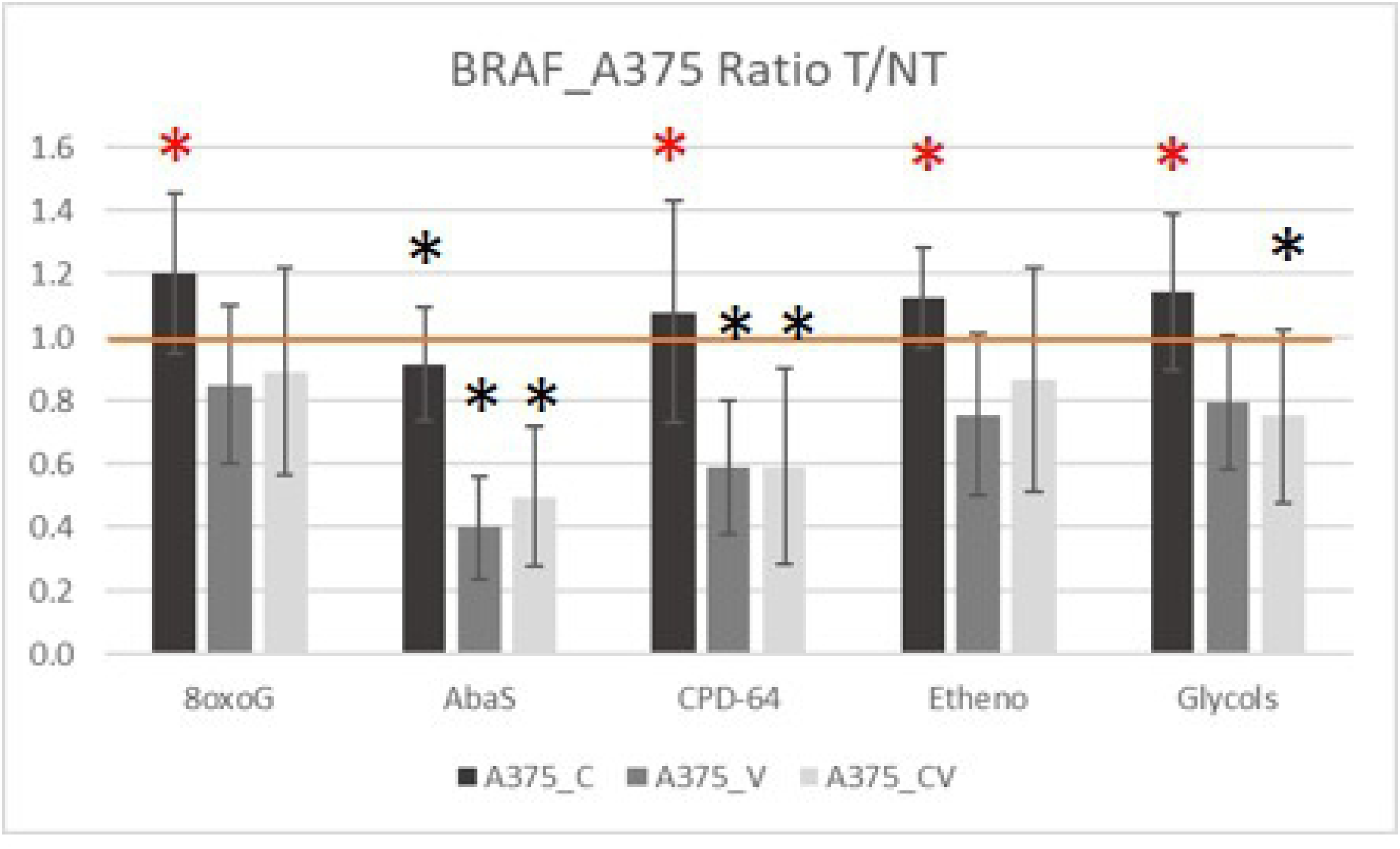
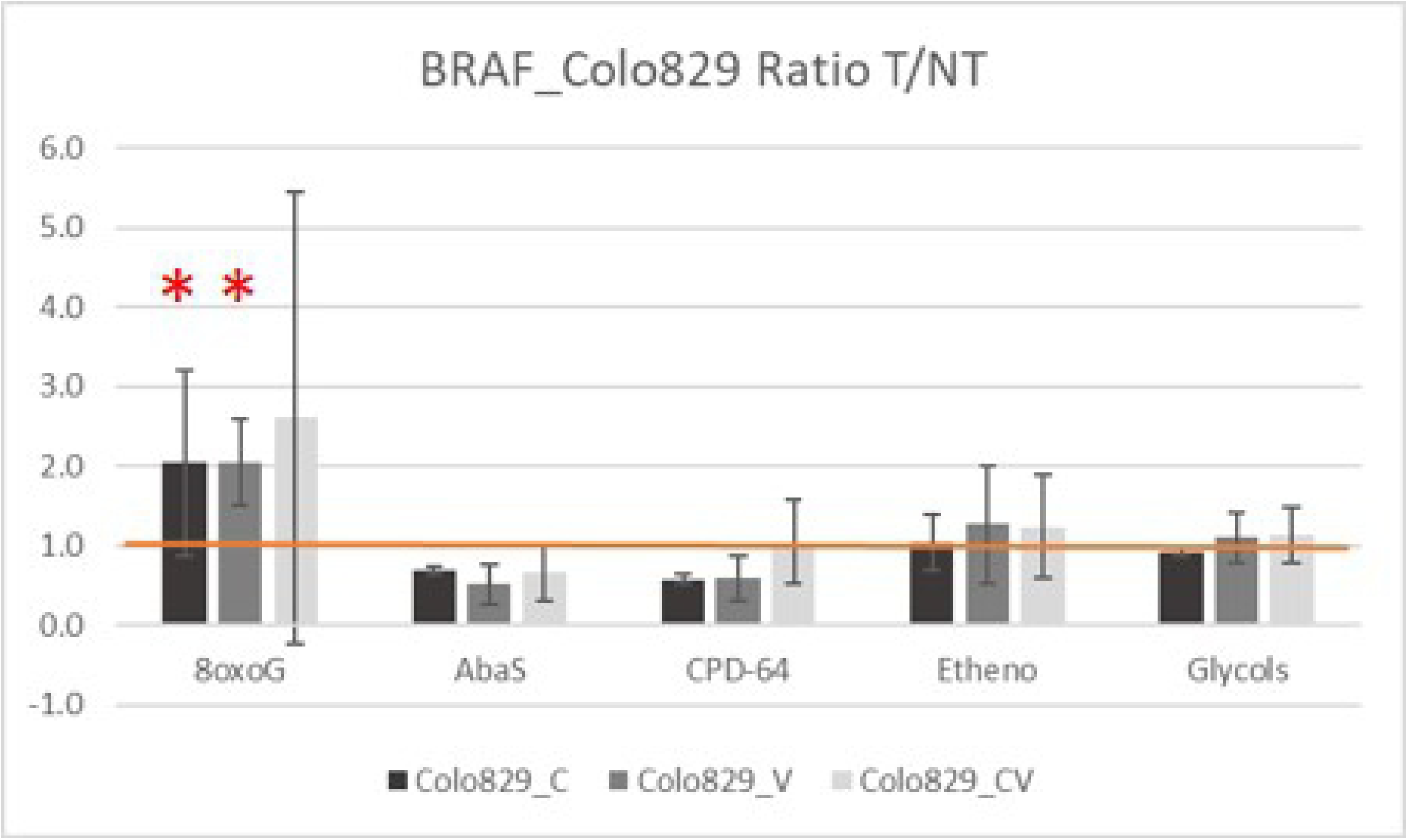
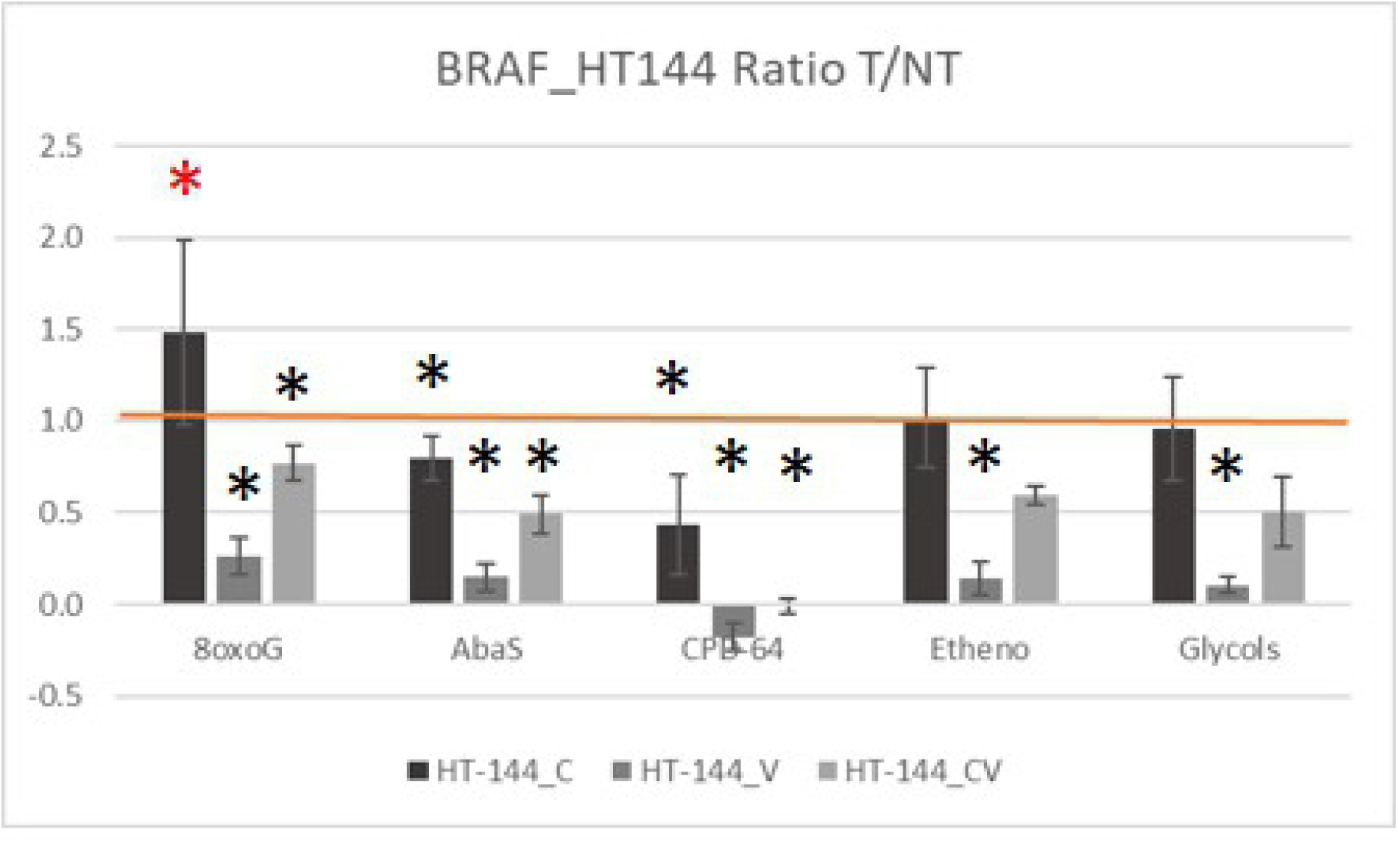
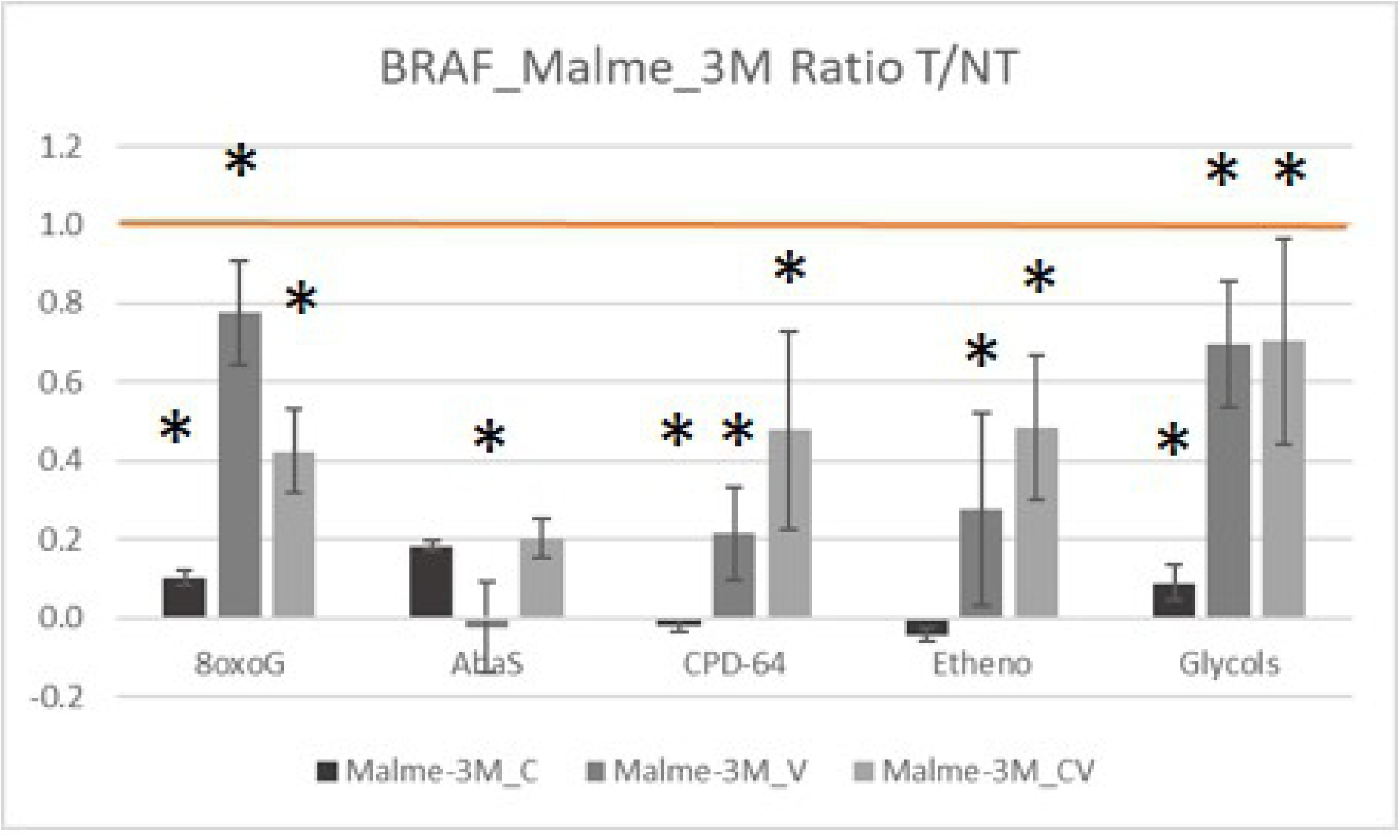
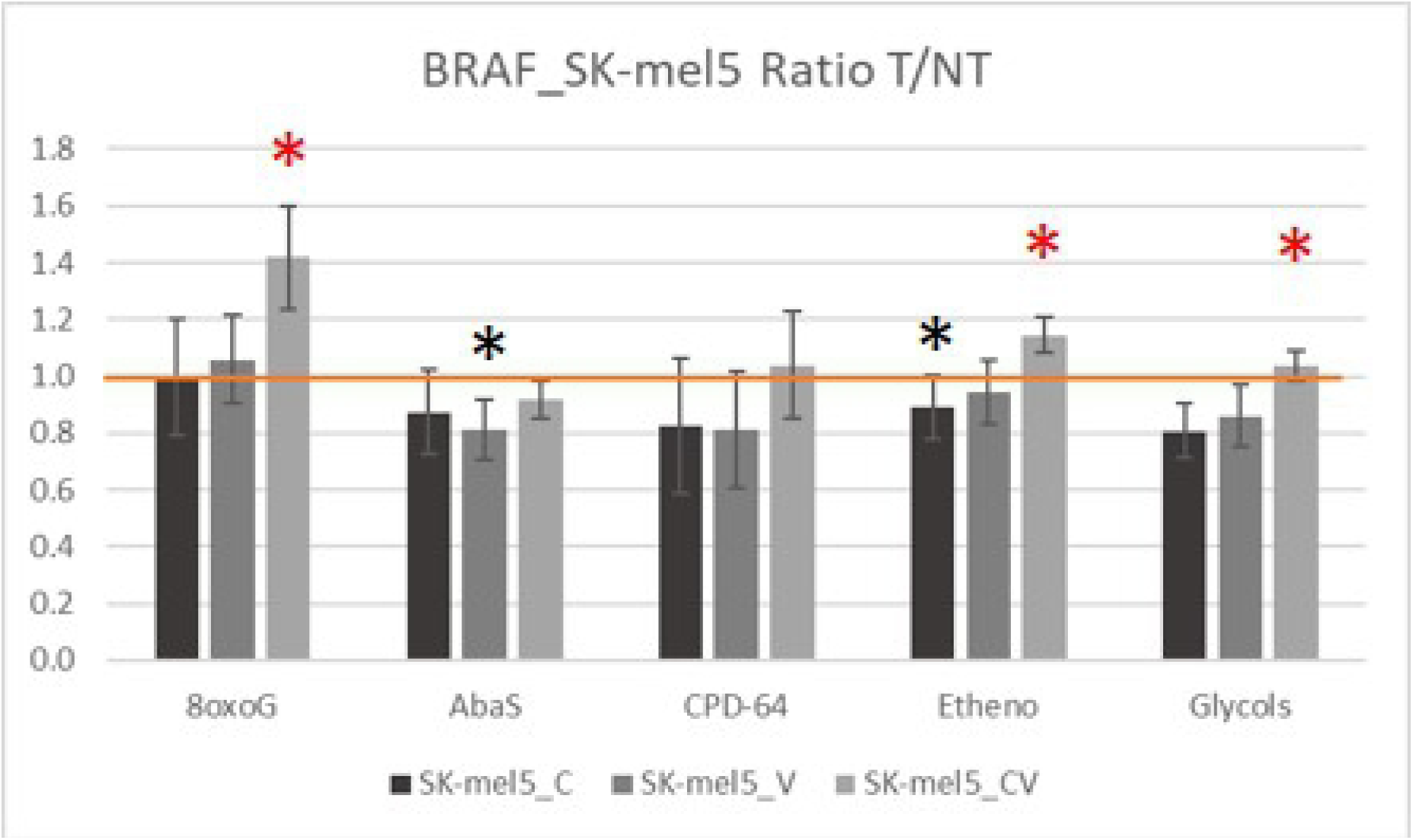
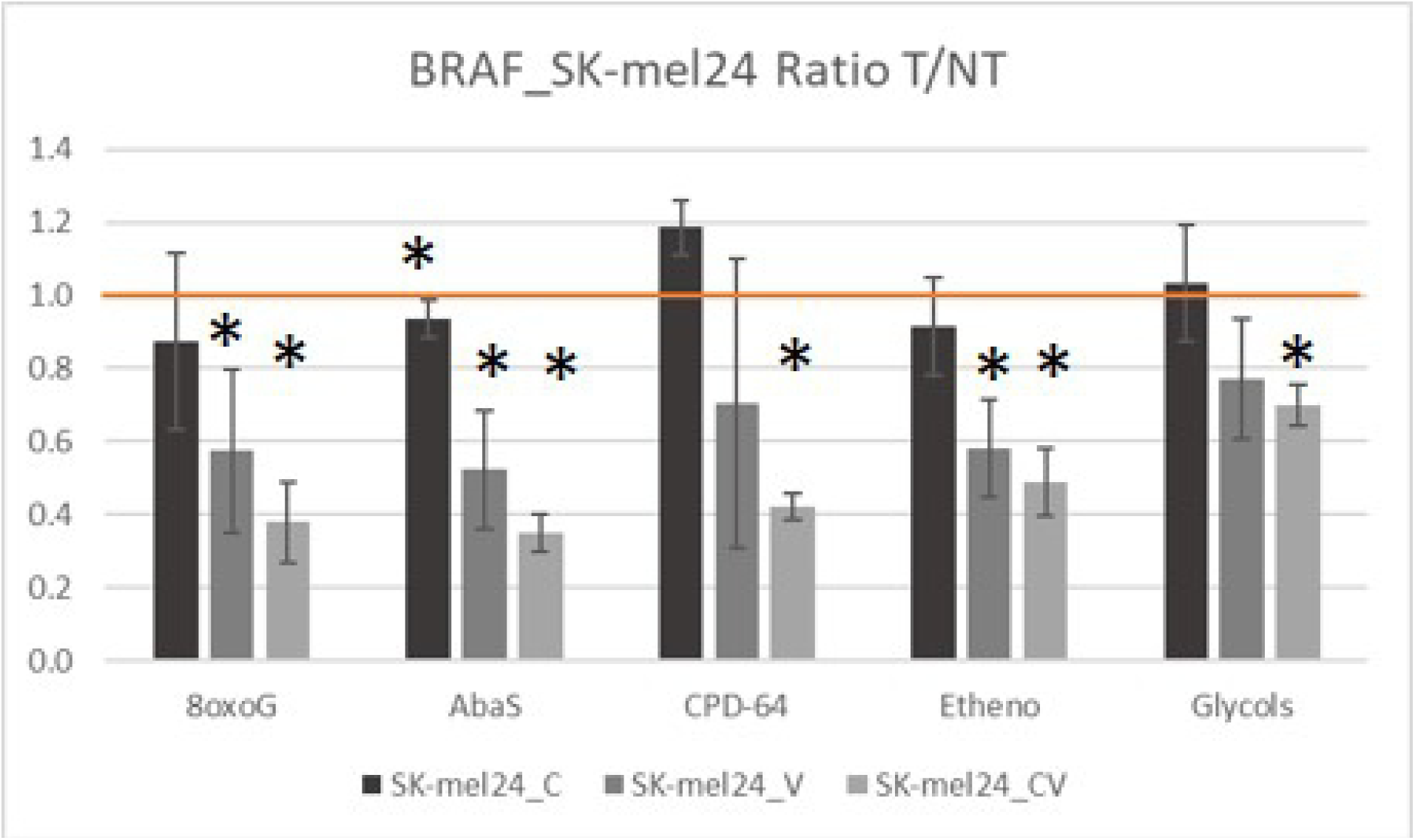
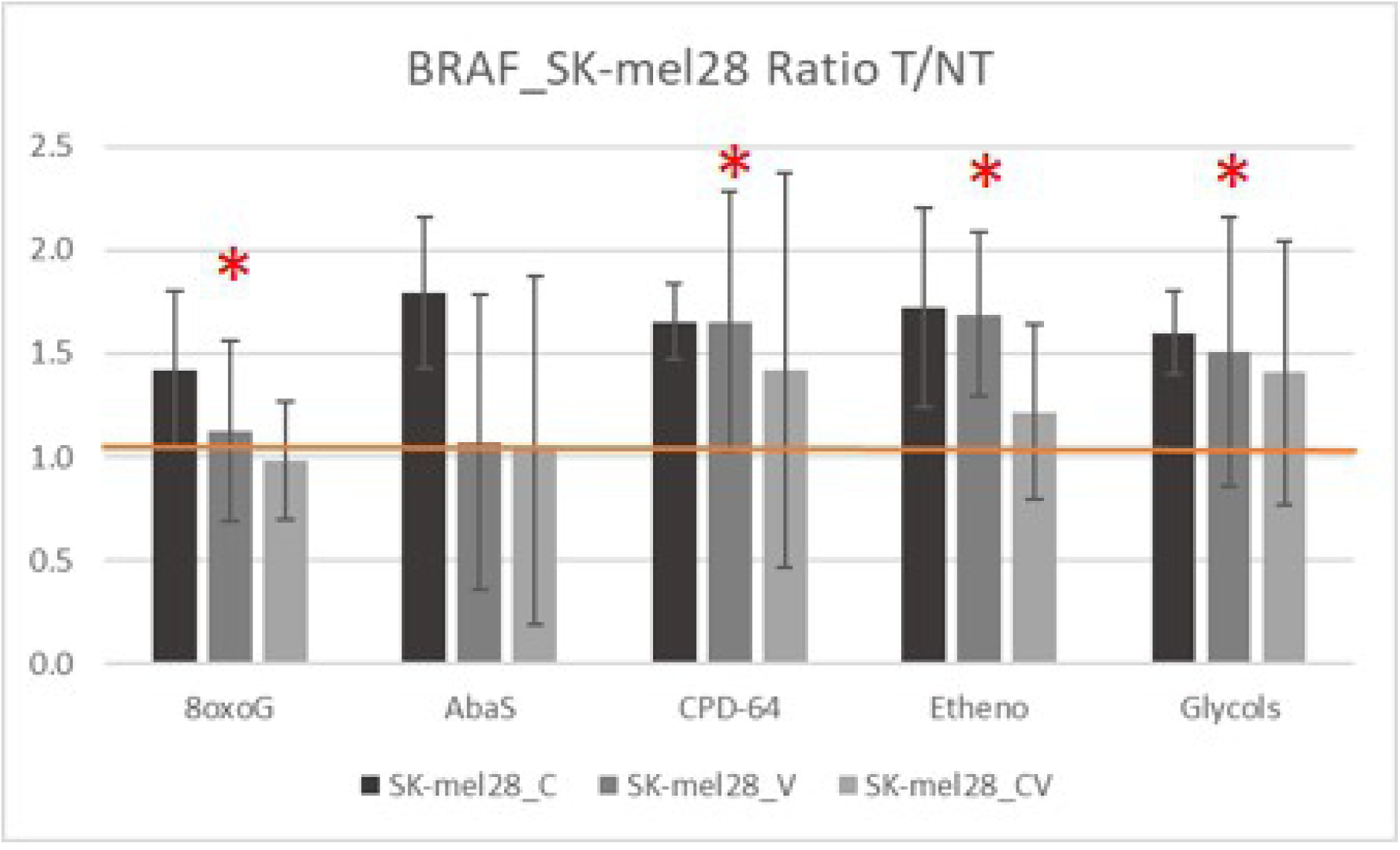
Effects of therapeutic agent treatment on the DNA repair capacities of the different cell lines. Ratio of the FI T/FI NT were calculated for the 3 biologically independent replicates. We used the student t-test to identify the results that were significantly different from 1, using the triplicated independent results. Cell lines and DNA repair substrates are designated. C= Cobi, V = Vemu, CV = combined treatment. *: p-value<0.05 compared to 1 (*inhibition of repair activity; * stimulation of repair activity)

As shown in Fig 3, treatment of the WT CHL-1 cell line with Vemu, and especially the drug combination, led to a significant stimulation of certain BER activities (repair of abasic sites, ethenobases and glycols). For the other WT cell line A7, no significant effect was observed, except an inhibition of the repair of abasic sites following Cobi+Vemu treatment. Different patterns were observed for the treated NRAS mutated cell lines. In particular, all repair BER activities (except repair of abasic sites) were significantly inhibited by Cobi in the case of MZ2 (Fig 3). For the other 2 NRAS cell lines, stimulation of repair of CPD-64, Etheno and Glycol were observed for M18 after Cobi treatment, whereas almost no change was detected for SK_mel2 after treatment. The DNA repair activities of the 3 BRAF mutated cell lines (HT-144, Malme-3M, SK-mel24) were strongly inhibited by Vemu or Cobi+Vemu. That was also true but to a lesser extent for A375. Negative impact of Vemu on SK-mel5 DNA repair activities was very moderate but some stimulation of several DNA repair activities following Cobi+Vemu treatment was observed for this BRAF mutated cell line. Unexpectedly Malme-3M was particularly sensitive to Cobi treatment with regards to DNA repair capacities. A375 or HT-144 treated with Cobi displayed a significant increase in the repair of abasic sites. The BRAF mutated cell line SK-mel28 also showed some stimulation of several DNA repair activities following Vemu treatment (Fig 3).

## Discussion

Melanoma can be a devastating form of skin cancer, and current treatment modalities are often ineffective. Thus, to better stratify melanoma patients that would exhibit a positive treatment response, improved diagnostic markers are needed. Given the important role that DNA repair plays in the etiology of cancer, including of the skin, we determined the DNA repair activities of several relevant cell lines using a precise tool that measures in parallel several excision/synthesis repair mechanisms, including defenses against oxidative DNA damage and DNA photoproducts. This tool has already been described in detail [25] and used to demonstrate that (i) chronic sun exposure impacts specifically the repair of DNA photoproducts [24] and (ii) DNA-damaging agents trigger specific DNA damage responses according to their mechanisms of action [29]. Our analysis herein shows that mutations in the BRAF or NRAS gene, major genetic drivers of melanoma, can affect the functional DNA Repair profiles of melanoma cells, with the excision/synthesis repair mechanisms of both BER and NER being influenced.

Globally, at basal level (i.e., untreated or NT), the NRAS mutated group displayed the highest DNA repair capacities as measured by the excision/synthesis repair assay, although it only reached significance for the repair of abasic sites compared to the WT group. Since oncogenic Ras induces genotoxic stress through the production of reactive oxygen species [30], and consequent increases in oxidative DNA damage [31], this may explain the observed increased repair in NRAS mutated cells. In addition, oncogenic Ras is frequently associated with activation of the DNA Damage Response (DDR), with the ATR protein kinase pathway being a key mediator [32]. Constitutively active NRAS protein also activates the MAPK and phosphatidylinositol 3-kinase (PI3K)/protein kinase B (AKT) signaling pathways [33], possibly contributing to the hyperactivation of DNA repair mechanisms in NRAS mutated cell lines. In contrast to NRAS mutation, the oncogenic BRAF mutation, which activates the MAPK pathway, was associated with decreased DNA repair activities. The connection between the MAPK pathway and the DDR, however, still requires further clarification, in particular to decipher which precise molecular mechanisms are associated with that decrease. At basal level, the repair of DNA photoproducts was reduced in the BRAF mutated group compared to WT and NRAS mutated groups, although statistical analysis did not reveal significance. The specific impact of mutation in the BRAF gene on the repair of DNA photoproducts, because BRAF mutations and UV-induced lesions are both hallmarks of melanoma, requires further investigation on a larger panel of cells.

Cobi alone affected the repair capacity of only one NRAS mutated cell line (MZ2), which changed from a high repair group to a low repair group after treatment. That feature was not observed when the Cobi+Vemu combination was used. Since Cobi is a selective inhibitor of MEK1 [34], it is tempting to speculate that pathway deactivation adversely impacts DNA repair capacity. However, it remains unclear why only one NRAS cell line was affected. More surprisingly, we observed that the WT CHL-1 cell line was sensitive to Cobi (S1 Fig). At basal level, this cell line had a DNA repair profile that matched the BRAF mutated “low repair” group, suggesting an inherently poor repair capacity. Treatment of CHL-1 cells with Cobi led to a further decrease in repair activities (significant for repair of 8oxoG, Fig 3), whereas treatment with Cobi+Vemu strongly stimulated some repair activities, suggesting paradoxical regulation of repair capacities. It would be interesting to characterize the MAPK pathway for CHL-1 cells, which behaved in an unexpected way.

In most cases, Vemu alone and Cobi+Vemu had similar effects. In particular, a global shift was noticed under these treatment conditions toward lower DNA repair capacities for BRAF mutated cell lines only, resulting in significant differences with the repair capacities of NRAS mutated cells (Figs 1-2). This observation is interesting as induction of secondary malignancies is a well-known side effect of BRAF inhibitors, such as Vemu [35], and a low level of DNA repair is associated with increased risk of developing cancer. The mechanism of action of Vemu involves selective inhibition of the mutated BRAF V600E kinase. The consequence is a reduced signaling through the aberrant MAPK pathway and the downstream kinases MEK1/2 and ERK1/2 [36]. In addition, it has been demonstrated that in keratinocytes, Vemu impedes the repair of ultraviolet A-induced photoproducts [37]. We showed here that Vemu not only affects the repair of DNA photoproducts by NER, but all BER activities as well, probably through its action on upstream regulation pathways. Another consequence of the treatment of cells with Vemu or Cobi+Vemu, because MAPK blockade with Vemu is mutation specific, was the clustering of all NRAS mutated cell lines, where DNA repair was not modified, in the same “high repair” group (Fig 2C,D).

At the basal level, three subtypes of BRAF mutated cells could be distinguished: (i) low repair, comprised of Colo-829, A375 and SK-mel24; (ii) an intermediate group that included SK-mel5, HT-144, Malme-3M; (iii) and high repair, consisting of SK-mel28. After treatment with Vemu and Cobi+Vemu, SK-mel28 and SK-mel5 were part of the high repair group. Notably, the WT CHL-1 cell line, which was unexpectedly sensitive to Cobi, displayed a strong stimulation of almost all DNA repair pathways when treated by Cobi+Vemu, supporting some signaling pathway dysregulation.

Collectively, our work found that certain cell lines showed an unexpected induction of either all repair pathways (as for SK-mel28) or only certain activities with specific treatment (Fig 3). This stimulation of DNA repair following treatment could be associated to some paradoxical activation of signaling pathways. Since mechanisms of resistance to Vemu are complex and multiple, they might involve upregulation of other components in the MAPK pathway [38] that can be revealed by a functional DNA repair assay.

In closing, different patterns of responses were observed by BRAF mutated and NRAS mutated melanoma cells lines before and after treatment with the therapeutic agents, Cobi and Vemu, either alone or in combination. The variability of cellular responses reported herein could reflect the heterogeneity in drug response observed in clinics following treatment with the targeted therapies. Our studies have also revealed the complexity of the signaling mechanisms that regulate DNA repair activities, and uncovered a role for the MAPK signaling cascade in the control of both BER and NER. Looking forward, a strategy of classification and profiling that involves a comprehensive analysis of DNA repair capacities could benefit the understanding of drug efficacy and drug resistance in melanoma. This is specifically important in the perspective, for example, of combining DNA damaging drugs with BRAF inhibition, which recently demonstrated synergies in their therapeutic potential [39].

## Acknowledgement

We thank Dr. David M Wilson III (Boost Scientific) for constructive input and manuscript editing.

The study was funded by Institut Roche, France.

## Supporting information captions

**S1 Table. Cytotoxicity experiments**

Concentration range of the drugs that were tested (Cobi alone, Vemu alone, and combination) for each mutation group (WT, BRAF, NRAS).

**S1 Fig. Cytotoxicity curves obtained for all cell lines.**

**S2 Table. Description of the data (FI) – Impact of the treatment by cell line.**

Description of each cluster by the variables (repair pathways): the v-test identified the variables that characterize each cluster (Noted Class_N in Fig 2). This test measured the deviation of the mean of the variable within a cluster, from the overall mean of the all set of data. A high absolute value of the v-test for the variables (repair pathway), indicated that the variable strongly contributed to the definition of the cluster.

Significance was reported as the p-value for each repair pathway.

